# Psychological stress and social support are associated with opposing single-cell pro-inflammatory gene regulatory mechanisms in adults

**DOI:** 10.1101/2025.10.14.682318

**Authors:** Ali Ranjbaran, Cindy Kalita, Julong Wei, Kristin M. Davis, Julian Bruinsma, Henriette Mair-Meijers, Gabrielle Garlicki, Adnan Alazizi, Michael Petriello, Yanping Jiang, Samuele Zilioli, Roger Pique-Regi, Francesca Luca

## Abstract

Psychological stress is linked to elevated markers of chronic inflammation, whereas social support is associated with lower levels; yet, the molecular mechanisms mediating these effects are poorly understood. We investigated gene regulatory variation in peripheral blood mononuclear cells (PBMCs) from 165 self-reported African American adults (aged 50-89 years) using single-cell RNA sequencing (scRNA-seq) and single-cell chromatin accessibility (scATAC-seq). Self-reported psychological stress and social support were associated with differential expression of 1,956 and 1,296 genes, respectively (10% FDR), primarily in CD4+ T cells and monocytes. Interferon signaling genes showed high expression in individuals with high psychological stress and low expression in those with high social support; this pattern mirrored gene expression in individuals with elevated circulating inflammatory markers (IFN-γ, TNF-α, IL-6). Genome-wide transcription factor (TF) motif analysis identified stress- and social support-associated changes in motif activity for 70 and 116 TFs, respectively, with 87 motifs enriched near differentially expressed genes. In CD4+ T cells, high psychological stress corresponded to increased IRF and STAT TF motif activity (interferon pathway), while social support was associated with reduced activity and expression in these pathways. We used an immune challenge paradigm (i.e., LPS stimulation), which confirmed the biological pathways of these gene regulatory effects. Our results demonstrate that psychological stress and social support modulate immune gene regulation at the single-cell level, revealing mechanistic links between psychosocial factors and inflammation, and suggesting that social support may promote immunological health.

## Introduction

Decades of research have established that the degree to which people perceive their lives as stressful and their social relationships as supportive contributes to health and disease. Robust evidence shows relationships between these two constructs—psychological stress and perceived social support—and various health outcomes, including the incidence of coronary heart disease(Richardson et al. 2012; Orth-Gomér, Rosengren, and Wilhelmsen 1993), asthma morbidity(Wisnivesky et al. 2010), and premature mortality(Keller et al. 2012; Shor, Roelfs, and Yogev 2013). Psychological stress, the subjective experience of feeling threatened in response to objective stressors, activates physiological responses (endocrine and immune) and prompts behaviors that can increase vulnerability to disease over time(McDade, Hawkley, and Cacioppo 2006; Knight et al. 2021; Fields et al. 2014). Similarly, perceived social support, defined as the perception that others are available to provide help when needed, has been found to be associated with reduced medical morbidity and premature mortality(Uchino 2009) through several pathways, including modulation of the immune system(Uchino et al. 2018). Importantly, psychological stress and perceived social support may affect biological mechanisms implicated in health and disease above and beyond objective exposure to stressful life events and levels of social support(Uchino 2009; Keller et al. 2012). Despite their important role in health and disease, a systematic understanding of how psychological stress and social support affect gene expression and its regulatory mechanisms has yet to be established.

The extant social genomic literature has revealed that a host of psychosocial factors are associated with a pattern of gene expression characterized by an upregulation of IL-6, TNF-α, and NF-κB-regulated genes and a downregulation of genes involved in antiviral responses and antibody synthesis(Steven W. Cole 2019). This pro-inflammatory transcriptional shift has been found in relation to acute psychological stress(Kuebler et al. 2015), loneliness(Steve W. Cole et al. 2007), socioeconomic disadvantage(Miller et al. 2009), and depression(Chiang et al. 2019). Two limitations of these pioneer studies include their focus on pre-specified gene sets (vs genome-wide gene expression) and reliance on bulk gene expression (vs. single-cell gene expression). Recent studies have addressed some of these limitations. For example, findings from the Multi-Ethnic Study of Atherosclerosis (MESA) specifically focused on monocyte gene expression and confirmed that psychosocial stressors, including psychological stress, were associated with expression of a pre-selected set of genes related to chronic inflammation(Brown et al. 2020). How psychological stress affects genome-wide gene expression in monocytes as well as other cell types remains to be established.

One important gap in the social genomic literature is the limited attention devoted to the regulatory mechanisms through which psychosocial factors influence gene expression. Transcription factors (TF), which modulate gene expression by binding specific DNA motifs, are key regulatory factors that may link psychosocial experiences to transcriptional changes. So far, social genomics studies have used sequence-based predictions of TF motif binding(Steve W. Cole et al. 2005). This approach has revealed enrichment of pro-inflammatory TFs binding motifs, such as NF-κB, nearby genes differentially expressed in association with several psychosocial factors, including loneliness, subjective social status, and neighborhood adversity(Steve W. Cole et al. 2007; Murray et al. 2019; Miller et al. 2022). One limitation of this approach is that it infers regulatory activity indirectly from downstream transcriptional changes and motif sequence rather than directly measuring chromatin accessibility. As a result, it cannot distinguish between functionally relevant accessible and occupied regulatory elements or identify chromatin dynamics. In contrast, ATAC-seq enables direct, genome-wide profiling of open chromatin regions, providing a more accurate and mechanistic view of TF motif activity and gene regulatory potential across immune cell populations. Interestingly, a recent study used ATAC-seq to profile chromatin accessibility in monocytes isolated from the bone marrow of mice exposed to chronic stress, showing higher chromatin accessibility at promoter regions of genes linked to IL-1 and IL-6 production, consistent with their higher gene expression levels. The study also demonstrated that monocytes from individuals with high psychological stress exhibited elevated NF-κB and interferon signalling, mirroring the gene expression patterns observed in the mouse model (Barrett et al. 2021). Another study has used ATAC-seq in immune cells from rhesus macaques of differing social status rankings and found that low social status macaques had significantly higher chromatin accessibility at promoter and enhancer regions of NF-κB, while the high-status macaques had higher accessibility for glucocorticoid response-associated transcription factors, which are involved in anti-inflammatory activity and NF-κB repression(Snyder-Mackler et al. 2019). These previous studies have either focused solely on myeloid cells or relied on bulk transcriptomic and epigenomic profiling, which masks the cellular heterogeneity of the immune cells and offers limited insight into responses in each immune cell type or the upstream regulatory mechanisms involved. Understanding how social factors shape the transcriptional landscape of immune cells is essential for identifying regulatory pathways that link the psychosocial factors to biological processes implicated in disease risk.

In the current study, we collected data on psychological stress and perceived social support levels in a sample of urban-dwelling middle-aged and older self-reported African American adults and performed single-cell RNA sequencing (scRNA-seq) and single-cell ATAC sequencing (scATAC-seq) in peripheral blood mononuclear cells (PBMCs) (Figure 1A). By integrating scRNA-seq and scATAC-seq, we directly assessed how psychological stress and perceived social support (hereafter, social support) influence the immune system at the gene regulatory level, and confirmed the biological significance of these gene regulatory effects following an immune challenge (i.e. LPS stimulation).

**Figure 1.**
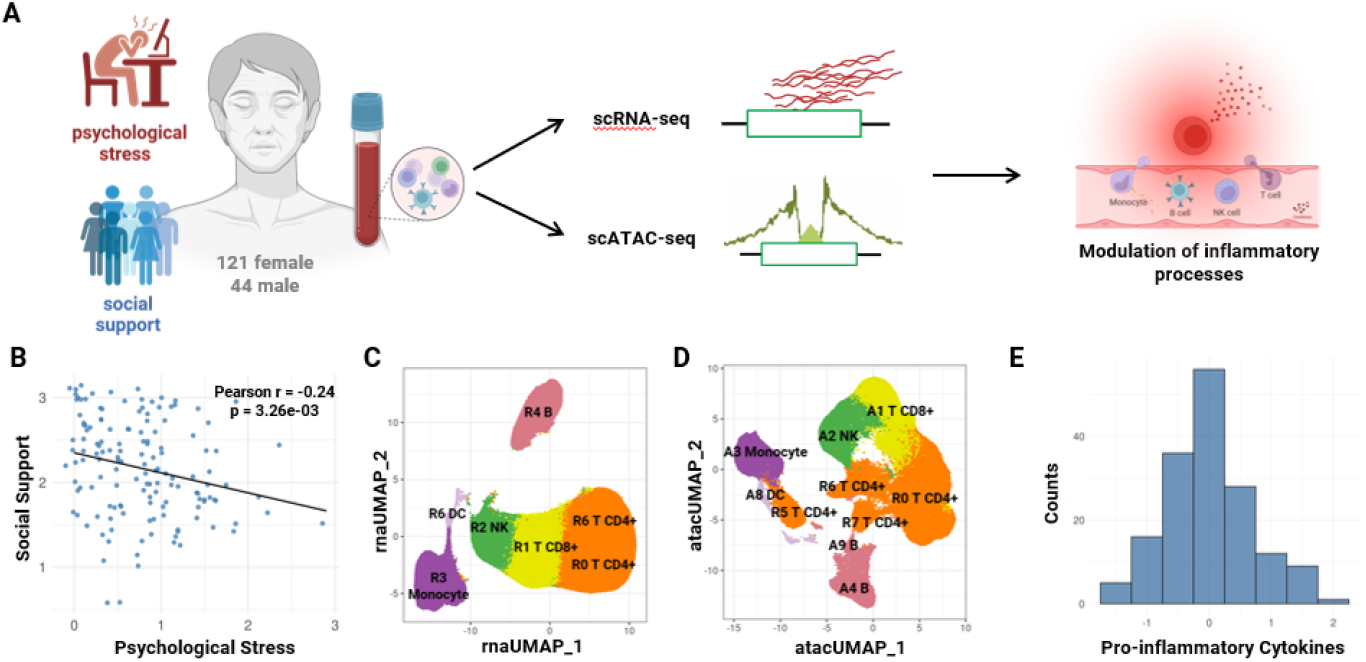
Study overview. (A) Levels of perceived psychological stress and social support were calculated in 165 self-reported African American adults aged 50-89 years old. PBMCs were collected, followed by genotyping, single-cell RNA sequencing, and single-cell ATAC sequencing. (B) Scatter plot of Pearson’s correlation of psychological stress and social support scores (C) UMAP visualization of scRNA-seq data for 8 clusters and 6 major cell types (D) UMAP visualization of scATAC-seq data for clusters for 9 clusters and 6 major cell types. (E) Histogram distribution of circulating pro-inflammatory cytokine composite calculated from IL-6, IFN-γ, and TNF-α.

## Results

### Psychological stress and social support influence the transcriptome in opposite directions

We measured single cell gene expression (RNA-seq) and chromatin accessibility (ATAC-seq) in PBMCs isolated from 165 adults participating in the HOLD (Health among Older adults Living in Detroit) study. HOLD is a study that investigates how biological, social, and psychological factors influence the health of urban middle-aged and older self-reported African American adults. We clustered 558,291 cells (RNA-seq) and 400,535 cells (ATAC-seq). Using a reference dataset(Ianevski, Giri, and Aittokallio 2022), we identified five major cell types from RNA-seq (Figure 1A), R0-CD4+ T cells (49.13%), R1-CD8+ T cells (18.90%), R2-NK cells (12.02%), R3-Monocytes (11.76%), and R4-B cells (7.92%) (Figure 1C; Figure S3, Table S3) and matched ATAC-seq cluster A0-CD4+ T cells (38.93%), A1-CD8+ T cells (19.30%), A2-NK cells (16.18%), A3-Monocytes (11.31%), A4-B cells (6.55%), A5-CD4+ T cells (2.45%), A6-CD4+ cells (2.41%), and A7-CD4+ T cells (1.87%) (Figure 1D; Figure S3, Table S4).

Our study focuses on two psychosocial variables: psychological stress and social support. Psychological stress is the degree to which a person appraises their recent life events and situations as uncontrollable and overloading. Social support is the degree of interpersonal resources a person perceives as available to support their needs. Considering these two variables separately, we found 1,956 differentially expressed genes (DEGs) associated with psychological stress across all cell types, predominantly in R3-Monocytes (1,366) and R0-CD4+ T cells (522) (Figure 2A). Social support was associated with a total of 1,296 DEGs across cell types, 457 in R0-CD4+ T cells, 411 in R1-CD8+ T cells, 164 in R2-NK cells, and 264 in R3-Monocytes (Figure 2A). In CD4+ T cells, psychological stress was primarily associated with higher gene expression, while in Monocytes, psychological stress was associated with both lower and higher gene expression. Social support was predominantly associated with lower gene expression.

**Figure 2.**
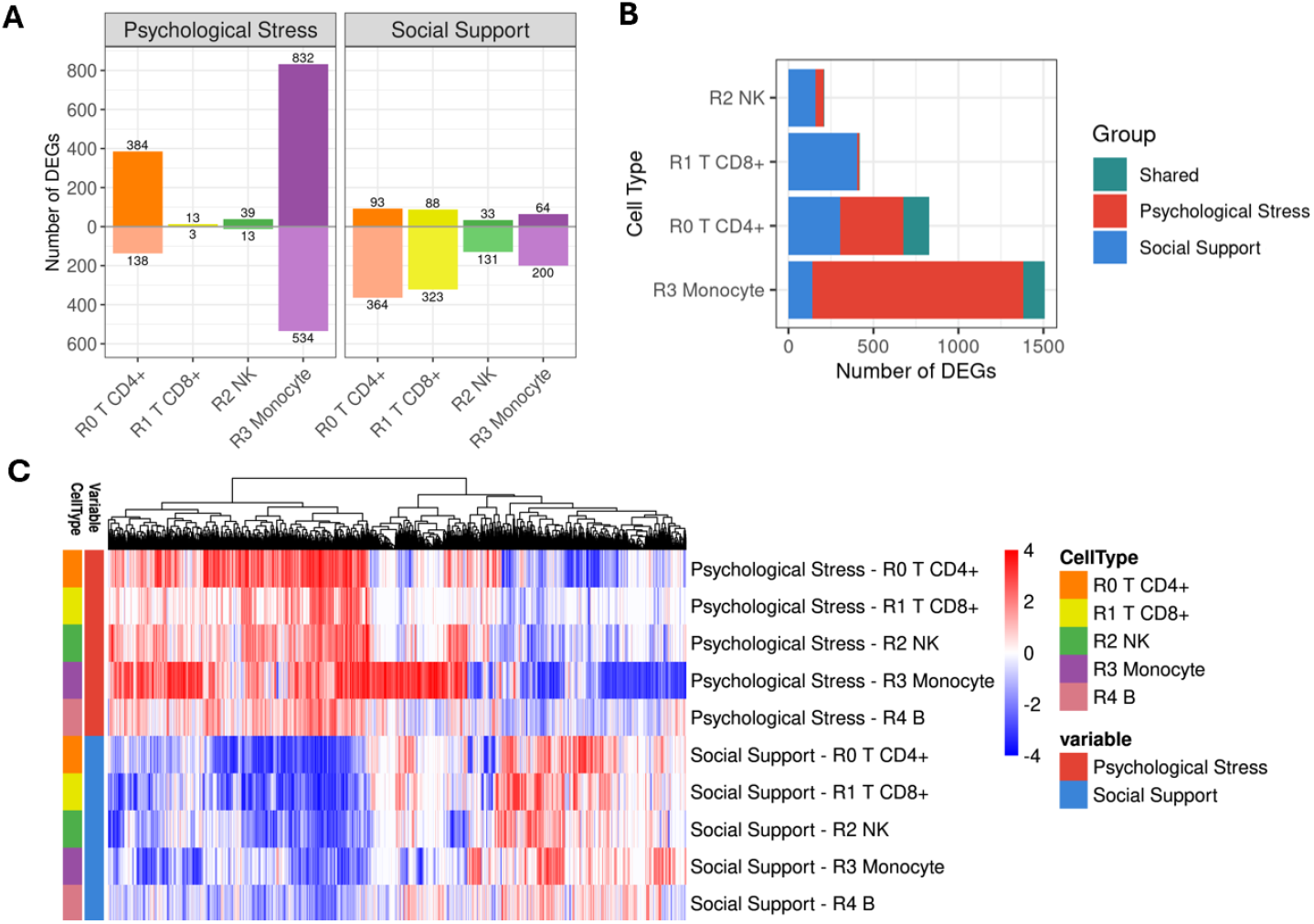
Identification of differentially expressed genes (DEGs) for psychological stress and social support. (A) Bar plot showing the number of up- and down-regulated genes per cell type at FDR 10%. (B) Number of shared and unique DEGs associated with psychological stress and social support for each cell type (C) Heatmap of z-score of 2,442 DEGs (columns) for psychological stress and social support across cell types (rows).

A closer look at the 2,442 unique DEGs identified across all cell types for either psychological stress, social support, or both revealed that genes with higher expression in individuals with elevated psychological stress tended to show lower expression in individuals with elevated social support (Figure 2C). This finding may partly be the consequence of the significant negative correlation between psychological stress and social support (r=-0.24, p-value=0.003; Figure 1B). To assess the unique effects of each of these variables on gene expression, we tested for DEGs associated with psychological stress while controlling for social support and vice versa. Out of the 1956 DEGs associated with psychological stress, 347 are DEGs after correcting for social support (17.7%), while 1,609 are not DEGs (82.3%). Additionally, we identified 9 new DEGs after correcting for social support. Of the 1,296 DEGs associated with social support, 91 are DEGs after correcting for psychological stress (7%), while 1,209 genes are not DEGs after correcting for psychological stress (93%) (Figure S11; Table S5-S6). While this may indicate a degree of confounding between the two variables, the number of genes identified did not have a lot of overlap (Figure 2B). Together, these results show that in this sample, it is difficult to separate the effects of psychological stress and social support on gene expression, as they likely acted with opposite effects in similar yet not identical pathways.

### Gene expression patterns associated with social support are only marginally explained by confounding factors

Socioeconomic status (SES) is a well-established social determinant of health(Braveman and Gottlieb 2014). To test the potential confounding role of SES on the gene expression pattern associated with psychological stress and social support, we used a differential gene expression model that controlled for SES. Out of the 1,956 DEGs associated with psychological stress, 795 are DEGs after correcting for SES (38.8%), while 1,197 are not DEGs (61.2%). Additionally, we identified 38 new DEGs after correcting for SES. These results indicate that the pro-inflammatory gene expression patterns associated with high psychological stress were mostly attributable to lower SES, but that psychological stress also exerted independent effects on transcriptional activity. Of the 1,296 DEGs associated with social support, 1,161 are DEGs after correcting for SES (89.6%), while only 135 genes are not DEGs after correcting for SES (10.4%). We also identified 35 new DEGs after correcting for SES. This finding suggests that the link between social support and decreased expression of pro-inflammatory genes was independent of SES (Figure S12; Table S7).

A second confounding variable we considered was body fat distribution as measured by waist-to-hip ratio (WHR). WHR is robustly associated with high levels of circulating markers of inflammation(Festa et al. 2001). To test if gene expression differences linked to psychological stress and social support were independent of body fat distribution, we considered a differential gene expression model controlling for WHR. Out of the 1,956 DEGs associated with psychological stress, we found 1,896 to be DEGs after correcting for WHR (96.9%), while only 60 of those were not DEGs after correcting for WHR (3.1%). Additionally, 32 new DEGs were associated with psychological stress after correcting for WHR. Out of the 1296 DEGs associated with social support, 1,238 were DEGs after correcting for WHR (95.5%), while 58 genes were not DEGs after correcting for WHR (4.5%). Additionally, we identified 88 new DEGs associated with social support after correcting for WHR. (Figure S13; Table S8). These results show that the gene expression patterns associated with psychological stress and social support are independent of body fat distribution.

### Gene expression patterns associated with social support are anticorrelated with those associated with pro-inflammatory cytokines

Pathways over-representation analysis of the genes with high expression in individuals reporting high psychological stress identified an enrichment of interferon signaling pathway genes, including IFN alpha/beta, interferon-gamma, and signaling by interleukins (Figure 2B). The same pathways were also enriched among genes with low expression in individuals with high social support. Given the observed enrichment of interferon pathway genes, we evaluated the relationship between gene expression levels associated with psychological stress and social support and markers of inflammation. Circulating levels of three proinflammatory cytokines, IL-6, TNF-α, and IFN-γ, were combined into a composite measure of systemic inflammation We compared the global gene expression differences associated with systemic inflammations with those associated with psychological stress and social support and found a positive correlation between systemic inflammation and psychological stress gene expression effects and a negative correlation between systemic inflammation and social support gene expression effects (Figure 3A). The strongest negative correlation was observed in Monocytes (r=-0.59), while the strongest positive correlation was observed in CD4+ T cells (r=0.43; Figure S10).

**Figure 3.**
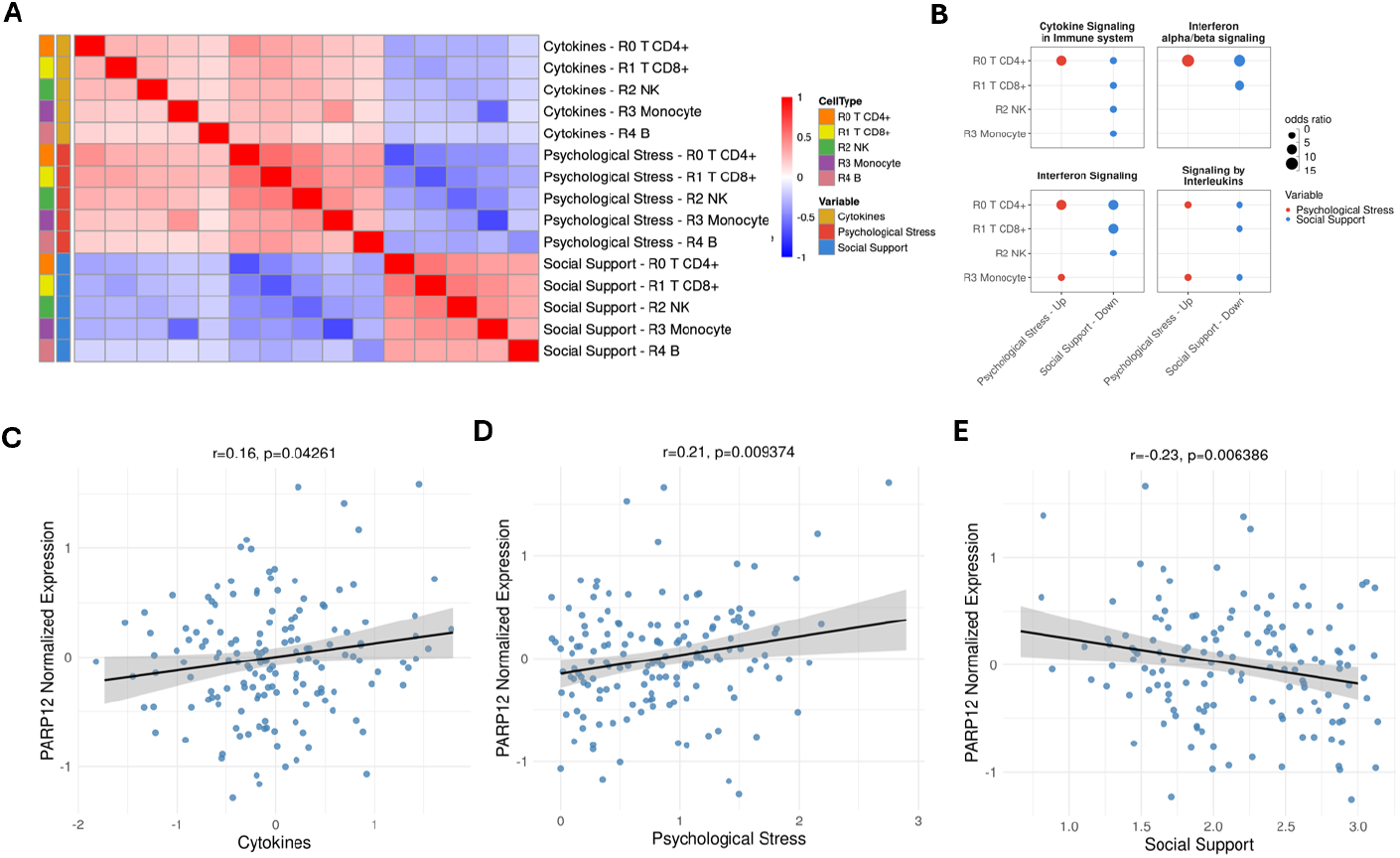
Correlation with pro-inflammatory signatures. (A) Heatmap of Pearson’s correlation of z-scores of cytokine gene expression effect with psychological stress and social support for each cell type. (B) Odds ratio of the pathway over-representation analysis across cell types for the top four most significantly enriched Reactome pathways in up-regulated DEGs in psychological stress and down-regulated DEGs in social support FDR 10%.(C-E) Scatter plot of the residual normalized expression of PARP12 gene in R3-Monocytes associated with cytokines, psychological stress and social support levels.

### Gene expression differences associated with psychological stress and social support are mediated by changes in transcription factor motif activities

We hypothesized that the gene expression differences associated with psychological stress and social support were mediated by differences in transcription factors (TF) binding (Figure 4A). To test this hypothesis, we derived the TF motif activities for each cell from scATAC-seq data, with high activity corresponding with strong binding. We identified a total of 158 Differentially Active Motifs (DAMs, 10% FDR, Figure 4B) associated with these factors across all immune cell types. Psychological stress was associated with 61 DAMs in CD4+ T cells, 6 in Monocytes, and 2 in CD8+ T cells. Social support was associated with 43 DAMs in Monocytes, 39 in CD4+ T cells, and 34 in NK cells. The effects on TF activity for psychological stress were negatively correlated with those for social support (Figure 4C), resulting in a pattern analogous to the one observed at the gene expression level (Figure 2C).

**Figure 4.**
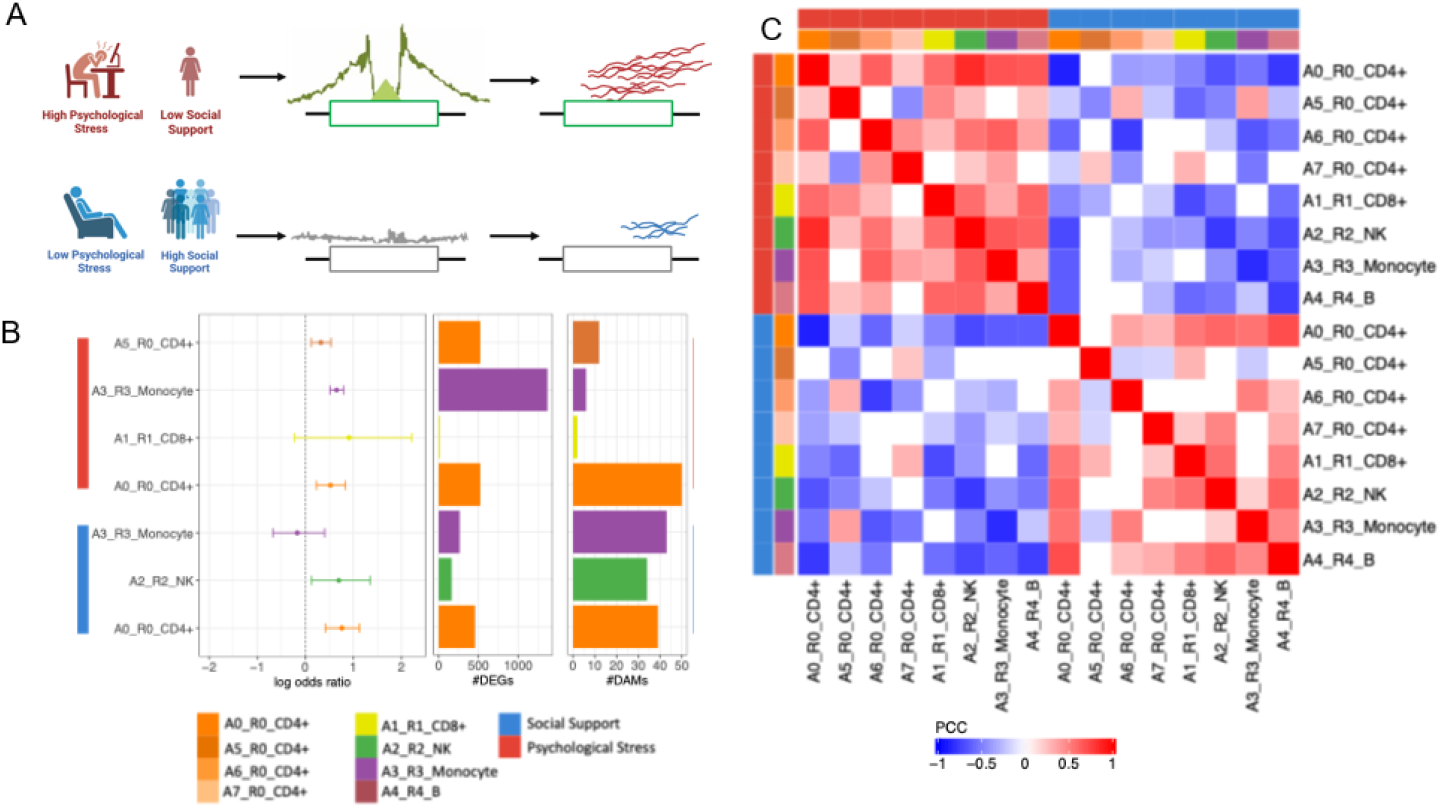
Differentially active motifs (DAMs) associated with psychological stress and social support. (A) Conceptual framework of the effect of psychological stress level on motif activity and gene expression (B) Forest plot visualizing log odds ratio (95% Confidence interval) of the enrichment of DEGs in DAMs in the corresponding cell types and variables, middle-bar showing the number of DEGs, and right-bar showing the number of DAMs. (C) Heatmap of Pearson’s correlation of z-scores for all tested motifs between psychological stress and social support across cell types. Colored by significance at p-value < 0.05.

We observed a significant enrichment of DEGs in DAMs for psychological stress in A0- and A5-CD4+ T cells (odds ratio-OR 1.684 and %95 CI 1.262-2.287, 1.380 and 95 % CI 1.125-1.701, respectively) and monocytes (OR 1.915, 95% 1.665-2.209) (Figure 4B). For social support, the DEGs in CD4+ and NK cells were enriched in DAMs in the corresponding cell types with odds ratios of 2.134 (95% CI 1.522-3.073) and 2.001 (95% CI, 1.131-3.858), respectively (Figure 4B). To distinguish what TFs were more impactful in modulating gene expression patterns associated with psychological stress and social support, we performed an enrichment analysis of DEGs for each individual TF. We identified 87 unique DAMs with significant enrichment (10% FDR) in DEGs across cell types and psychological stress and social support. Specifically, we observed that TFs that were key regulators of interferon signalling pathways (i.e., IRF2, IRF4, IRF9) showed high activity with psychological stress in CD4+ T cells and low activity with social support. Similarly, STAT1 showed high motif activity with psychological stress and low activity with social support in monocytes. (Figure 5A-B).

**Figure 5.**
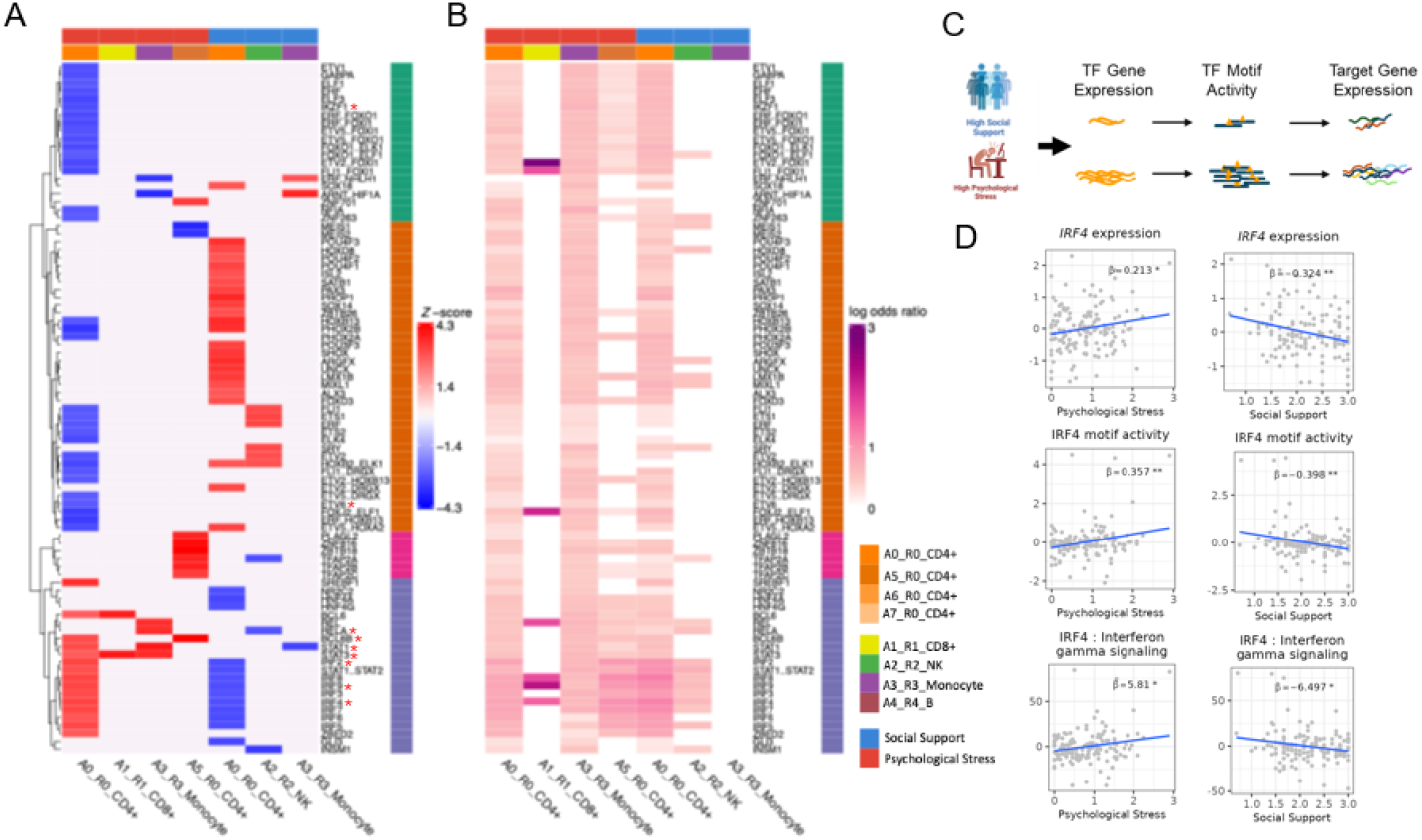
TF motifs in DEGs. Here, we defined a gene with the binding of TF to be that annotated in a peak containing a TF motif. Asterisks highlight DEGs. (A) Z score of TF activity associated with psychosocial factors (DAMs, 10% FDR). TFs clustering into 4 groups using a hierarchical method with complete linkage and Euclidean distance. Asterisk denotes TF genes that are DEGs in the corresponding cell type (B) Log odds ratio of the enrichment of individual TF motifs in DEGs in the corresponding cell type and psychosocial variables. (C) schematic overview of the underlying mechanism of TF genes driving target gene expression changes (D) IRF4 gene expression (to row), motif activity (middle row), and pathway score of the IFN--γ signaling pathways belong to the IRF4 TF (bottom row) associated with psychological stress levels (left) and social support (right).

From the 87 unique DAMs with significant enrichment, we identified 9 unique TF genes to be differentially expressed in the corresponding cell type (Figure 5A). These factors belong to the STAT, IRF, and BCL family of TF, which serve as key master regulators of numerous inflammatory functions in immune cells, particularly CD4+ T cells. For all these TF, high psychological stress was associated with high expression and motif activity of the TF, while high social support was associated with low expression and activity (Figure S16). These results suggest that psychological stress and social support influence TF gene expression, leading to changes in TF motif activity that ultimately affect the downstream pathways these factors regulate (Figure 5C). For example, we observed that individuals with high psychological stress had high expression of the IRF4 gene, high TF motif activity at the IRF motif, and elevated IRF4 reactome pathways scores, including interferon-γ signaling pathways in CD4+ T cells, while the opposite was observed for individuals with high social support (Figure 5C-D).

### Genes with high expression in individuals experiencing high psychological stress and low social support are up-regulated following LPS stimulation

To confirm the pro-inflammatory function of genes highly expressed in unstimulated cells from individuals with high psychological stress and low social support, we stimulated PBMCs in parallel with lipopolysaccharide (LPS), a bacterial cell membrane component widely used to simulate bacterial infection and inflammation, and performed scRNA-seq and scATAC-seq. We observed a predominantly concordant pattern between response to LPS and genes associated with psychological stress, with 743 genes that had high expression in individuals with high psychological stress having increased expression in response to LPS. Similarly, 442 genes that had low expression in individuals with high psychological stress had lower expression in response to LPS. We only observed 82 genes that had effects in the opposite direction in response to LPS and as a function of psychological stress. As expected, social support-associated genes had a discordant effect in response to LPS, with 648 that had low expression in individuals reporting high social support having increased expression in response to LPS. Similarly, 137 genes with high expression in high social support individuals had lower expression under LPS treatment. We only observed 33 genes with effects in the same direction in response to LPS and as a function of social support (Figure 6A; Table S9). We observed that this pattern was most prominent in CD4+ T cells and Monocytes (Figure 6C; Table S9).

**Figure 6.**
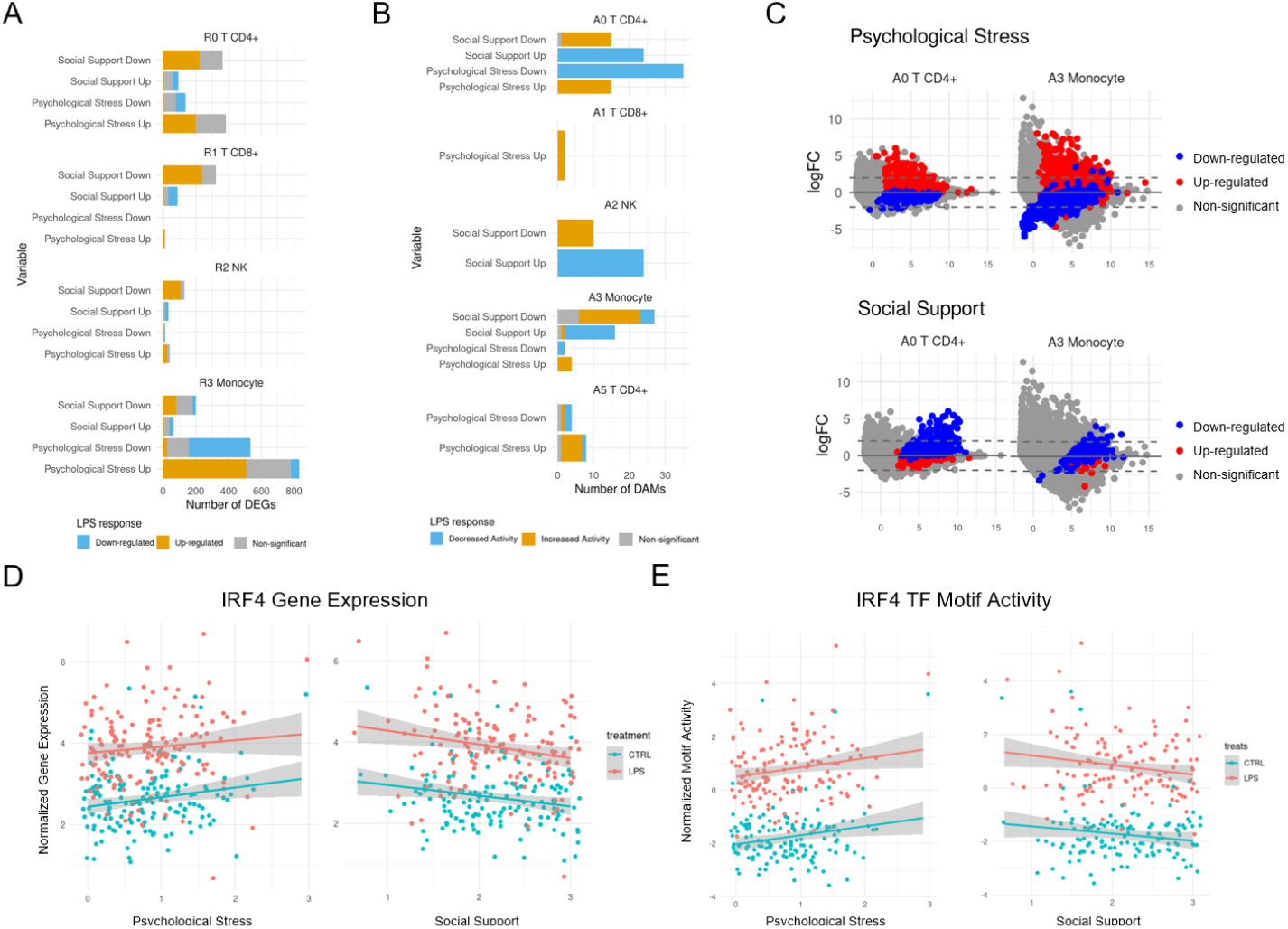
Psychological stress and social support associated DEGs in LPS-treated cells (A) Stacked bar plot of the number of LPS response DEGs separated by up- and down-regulated genes associated with psychological stress and social support per cell type. (B) Stacked bar plot of the number of LPS response DAMs separated by increased or decreased motif activity in association with psychological stress and social support, per cell type. (C) MA plot of LPS vs CTRL treatment colored by up (red) and down (blue) regulated genes associated with psychological stress (top) and social support (bottom). (D) Normalized expression and (E) TF motif activity of IRF4 in R0 A0 CD4+ T cell in LPS and CTRL treatment correlated with psychosocial stress and social support.

Using differential motif activity analysis in response to LPS, we observed a similar pattern in the gene regulatory mechanism to the one observed in the case of gene expression. Twenty-seven motifs that had high activity in individuals with high psychological stress also had high activity in response to LPS. Thirty-nine DAMs with low activity in individuals with high psychological stress had low activity in response to LPS. We only identified 2 DAMs with opposite effects between response to LPS and in association with psychological stress. When we considered the DAMs associated with social support, we observed that 41 of the motifs that had low activity in high social support individuals had a high activity in response to LPS. Similarly, 62 motifs with high activity in individuals with high social support had a low motif activity in response to LPS. Only 5 TF motifs had effects in the same direction between the response to LPS and social support. Together, these results suggest that across all immune cell types considered, individuals with high psychological stress present activation of gene regulatory pathways that are also upregulated in response to LPS. Conversely, individuals with high social support present inactivation of pathways that are upregulated in response to LPS (Figure 6B; TableS10). For example, in CD4+ T cells, IRF4 gene expression was upregulated and its motif activity was increased in response to LPS. We observed that this TF had a high expression and motif activity in CD4+ T cells of individuals with high psychological stress and low expression and activity in individuals with high social support (Figure 6D-E).

## Discussion

In this study, we identified transcriptional and gene regulatory signatures in peripheral immune cells associated with psychological stress and social support in a sample of middle-aged and older self-reported African American adults. As people age, chronic low-grade inflammation becomes more prevalent and contributes to age-related conditions, such as cardiovascular diseases, making the immune regulatory state in aging adults a critical determinant of healthspan(Li et al. 2023; Zakai et al. 2007). Understanding how psychosocial factors such as psychological stress and social support influence immune regulation in aging adults is essential for uncovering the molecular mechanisms through which social experiences contribute to health outcomes(Valtorta et al. 2018; Redmond et al. 2013). Using single-cell genomics, we found that individuals reporting high levels of psychological stress displayed high expression of pro-inflammatory genes, including IFN response and antiviral defense pathway genes, while individuals reporting high social support exhibited low expression of pro-inflammatory genes and TF motif activity, particularly within CD4+ T cells. These findings suggest that psychological stress is associated with higher activation of the immune system, and, in contrast, social support is associated with lower immune activation and preserved homeostatic regulation of inflammatory responses in aging individuals.

Our findings pertaining to each cell type highlight the value of a single-cell approach in social genomics. We identified pronounced alterations in gene expression and motif activity within CD4+ and monocytes, two immune cell types central to acute and chronic inflammation, in association with psychological stress and social support. For example, we observed that in CD4+ T cells, genes related to IFN signaling, such as ISG15 and STAT1, had high expression in individuals with high psychological stress and low expression in individuals with high social support. Additionally, our chromatin accessibility analyses revealed differences in activity at key immune transcription factor motifs, including reduced activity of IRF and STAT family motifs in individuals with high social support, suggesting that positive social experiences shape regulatory dynamics upstream of pro-inflammatory pathways, thus reducing inflammation and potentially contributing to better health outcomes.

Importantly, we found that genes showing low expression in individuals with high social support tended to show high expression following LPS stimulation, whereas genes showing high expression in individuals with high psychological stress also exhibited up-regulation with LPS stimulation. Our findings extend previous work on chronic stress in model organisms(Barrett et al. 2021) to humans and further reveal, at a higher resolution, that both monocytes and CD4^+^ T cells are affected. Further, we demonstrated that these pro-inflammatory immune responses were inactive in individuals with high social support. We argue that improvements in a social support system may be helpful in overcoming some of the immunological challenges that have long been associated with chronic stress and adversity.

Importantly, these results were derived from a sample of urban-dwelling middle-aged and older African American adults, a demographic group that is often underrepresented in genomics research despite experiencing disproportionate exposure to stress and chronic health conditions(Hatch and Dohrenwend 2007; Aggarwal et al. 2021; Popejoy and Fullerton 2016; Wojcik et al. 2019). Our findings underscore the biological relevance of interpersonal resources as a potential modifiable factor in the immune aging process. Given that chronic low-grade inflammation is a known driver of age-related diseases, the observed anti-inflammatory signatures associated with social support may help explain epidemiological links between positive psychosocial factors and better health outcomes in later life.

This study also illustrates the power of integrating single-cell transcriptomic and epigenomic data to investigate the underlying molecular mechanisms of psychosocial experiences on human health. By resolving effects at the level of individual immune subtypes, we identified regulatory mechanisms that would be obscured in bulk analyses. The concordance between gene expression and transcription factor motif activity results further supports a coordinated shift in regulatory programs in response to the social environment.

While the dataset includes rich psychosocial phenotyping, the molecular data are cross-sectional, limiting assessment of longitudinal and chronic social stress and support on human health. Additionally, some immune cell subtypes were sparsely represented in the data or absent in PBMCs, limiting the scope of our study to the most abundant PBMC components.

In summary, our results support the hypothesis that social support is biologically embedded within immune regulatory mechanisms and suggest that improving interpersonal resources and overall social support may help mitigate pro-inflammatory immune states in older adults. Our study provides critical insight into the immune mechanisms through which these two established psychosocial determinants of health affect immunological health and contribute to disease vulnerability. Future work should examine these associations longitudinally and in larger populations, as well as assess the potential for interventions that enhance social support to reduce inflammation and improve healthspan.

## Methods

### Study participants

Participants are part of the Health among Older Adults Living in Detroit (HOLD) study, which investigates how biological, social, and psychological factors influence the health of middle-aged and older self-reported African American adults residing in Detroit, Michigan. Data collection for the study ran from November 2017 to March 2020. Of the 211 HOLD participants, we included in this study 165 (121 Females, 73%) participants who had available PBMCs and DNA samples, and measured psychosocial variables. The age of the selected participants was 50 to 89 years old (Figure S8C). The study was conducted in three parts: (1) an initial home visit, (2) a 5-day period during which participants completed questionnaires and study procedures following instructions they were given at the first home visit, and (3) a follow-up, second home visit. At the initial home visit, participants were given a detailed description of the study and the opportunity to ask questions prior to signing the consent form. During a 5-day period after the initial home visit, the participant would be responsible for completing daily diary questionnaires, providing saliva samples four times a day, logging their sleep via a wrist actigraph unit, and completing other questionnaires. On the second home visit, these materials were collected. Furthermore, blood pressure, heart rate variability, height, weight, waist-hip ratio, peak flow, and grip strength were also measured. At the end of the second home visit, a phlebotomist performed a blood draw.

### Psychosocial measures

Social support was measured using the 12-item version of the Interpersonal Support Evaluation List(Sheldon Cohen and Hoberman 1983; Sheldon Cohen and Wills 1985). This scale assesses perceived social support across three primary domains: appraisal, belonging, and tangible support. The scale includes a series of items about various aspects of social support (e.g., “There is someone I can turn to for advice about handling problems with my family,” “If I wanted to have lunch with someone, I could easily find someone to join me”). Response options range from 0 (definitely false) to 3 (definitely true). Negatively worded items were reverse-coded, such that higher values reflect greater social support. Overall scores were calculated by taking the mean of all items, then multiplying by the number of items (i.e., 12) to generate mean-imputed sum scores representing perceived social support with a possible range of 0-36. Individual questionnaire items can be found in Table S1-2.

Psychological stress was measured using a version of the four-item Perceived Stress Scale(S. Cohen, Kamarck, and Mermelstein 1983; Ohen and Williamson 1988) that was modified for use in our daily diary protocol (e.g.,”Thinking about today, how often did you feel that you were unable to control the important things of the day?”). Response options ranged from 0 (never) to 4 (very often). Positively-worded items were reverse-coded, such that higher scores indicated greater perceived stress. Participants were asked to complete the daily diary for five consecutive days. On days when all four items were answered, daily scores were calculated by taking the mean of all items. The average of all available daily scores were then taken to generate person-level psychological stress scores.

Socioeconomic Status (SES) was assessed using a composite score that combined participants’ self-reported household pre-tax income and self-reported highest educational attainment. Response options for income ranged from 1 (<$5000) to 13 (≥$150,000). Education response options ranged from 1 [No school/some grade school (1-6)] to 12 (Ph.D., Ed.D., MD, DDS, LLB, LLD, JD, or other professional degree). Income and education were each Z-scored, and the mean of the standardized values was taken to derive the SES composite score.

### Cytokine measures

#### Preprocessing

Blood samples were collected in 8.5mL serum separator tubes (BD Vacutainer SST, catalog no. 367988, BD, Franklin Lakes, NJ, USA) for assessment of circulating cytokines. Serum samples were cryopreserved in the Luca Lab and sent to the Petriello lab at Wayne State University to be assayed for circulating cytokines. The Meso Scale Diagnostics® (MSD) Multi-Spot Assay System (U-PLEX Proinflammatory Combo 1™; MSD Maryland, USA) was used, following manufacturer protocols. Manufacturer reported lower limits of detection (LLOD) are 1.7 pg/mL for IFN-γ, 0.14 pg/mL for IL-10, 0.69 pg/mL for IL-12p70, 3.1 pg/mL for IL-13, 0.15 pg/mL for IL-1β, 0.70 pg/mL for IL-2, 0.076 pg/mL for IL-4, 0.33 pg/mL for IL-6, 0.15 pg/mL for IL-8, and 0.51 pg/mL for TNF-α. Values below the LLOD for each cytokine were replaced with 0, in line with previous work(Van Bogart et al. 2021). More than 40% of blood samples were below the LLOD for several cytokines, including IL-12p70 (42.2%), IL-13 (56.8%), IL-1β (40.5%), IL-2 (73.5%), and IL-4 (69.7%). Due to their low detection rates, these cytokines were not included in analyses. Remaining cytokines (IFN-γ, IL-10, IL-6, IL-8, and TNF-α) were cleaned in line with existing research: each marker was log10(x+1) transformed; outliers > 3 SD from the mean were winsorized to ±3 SD of the mean(Graham-Engeland et al. 2018; Van Bogart et al. 2021).

#### Circulating Pro-inflammatory Cytokine Composite

To reduce the number of comparisons, the use of inflammatory composite scores has been recommended over analysis of individual cytokines(Fagundes et al. 2019). A cytokine composite representing circulating inflammation was derived using exploratory factor analysis (EFA(Van Bogart et al. 2021)). Each cytokine was Z-scored prior to EFA. Bartlett’s test of sphericity was significant (χ^2^ = 39.74, p <.001) suggesting that factor analysis was appropriate(Bartlett 1950). Given the present focus on pro-inflammatory physiology we conducted EFA on only the pro-inflammatory cytokines (i.e., IFN-γ, IL-6, IL-8, and TNF-α), excluding the anti-inflammatory IL-10. Both Eigenvalues and the scree plot indicated a one factor solution. In this solution, IFN-γ, IL-6, and TNF-α loaded onto the factor (loadings >0.40), while IL-8 did not. Thus, to create the circulating pro-inflammatory cytokine composite, the mean of z-scored IFN-γ, IL-6, and TNF-α was taken.

### Waist-to-Hip ratio

During the second home visit, waist and hip circumference (cm) were measured by trained research assistants using a standard tape measure. Waist-to-hip ratio was then calculated as waist circumference divided by hip circumference

### PBMC isolation and culture

PBMCs were extracted using a previously published Ficoll centrifugation protocol (Weckle et al. 2015), cryopreserved in freezing media, and stored in liquid nitrogen until the day of the experiment. All individuals in this study were genotyped from low-coverage (1×) whole-genome sequencing and imputed to 37.5 million variants using the 1000 Genomes database as reference and GLIMPSE2(Rubinacci et al. 2023). Cells were processed in batches of 12 donors on separate dates. For each batch, PBMCs were removed from liquid nitrogen storage and quickly thawed in a 37°C water bath before diluting with 6 ml of warm starvation media (90% RPMI 1640, 10% CS-FBS, 0.1% Gentamycin) and counting using Trypan blue staining on Invitrogen Countess II (Life Technologies Corporation, Bothell, WA). Cells were subsequently centrifuged at 400 xg for 10 minutes and resuspended in culture medium at 2 × 10^−6^cells/ml. 100μl of cell suspension was plated in each of 2 wells of a 96-well round-bottom cell culture plate for each sample. Cells were incubated in starvation media overnight (approx. 16 hours) at 37°C and 5% CO2.

### PBMC pooling and treatment

The following morning each of the 2 wells for each individual was treated with either: 10ng/ml LPS or vehicle control (water) for 6 hours. Cells were then pooled across individuals for a total of two treatment-specific pools. The pools were centrifuged at 300x g for 5 min at 4°C, washed with 5 ml ice-cold PBS+ 0.04% BSA, and centrifuged again. Each pool was resuspended in 500μl ice-cold PBS + 0.04% BSA and filtered through a 40μm Flow-Mi strainer (SP Scienceware, Warminster, PA). Cell concentration was determined using Trypan blue staining on Countess II and adjusted to 2 × 10^™6^cells/ml.

### scRNA-seq and scATAC-seq sample processing

Each pool was divided into 2 aliquots for scRNA-seq and scATAC-seq processing, respectively. For the scRNA, each pool was loaded onto a separate channel of the 10x Genomics Chromium instrument (10x Genomics, Pleasanton, CA), according to the manufacturer’s protocol, using v3.1 Dual index chemistry. Library preparation was done according to the manufacturer’s protocol. For scATAC-seq, cells underwent DNase treatment and nuclei isolation followed by library preparation according to the 10X Genomics protocol. Library preparation was performed according to the manufacturer’s protocol, using Single Cell ATAC kit V2.

### scRNA-seq raw data processing (alignment, demultiplexing, and cell type assignment)

The raw FASTQ files were aligned to the GRCh38.p12 human reference genome using CellRanger. We eliminated debris-contaminated droplets using the DIEM R package(Alvarez et al. 2020). The aligned count matrix was converted into a Seurat object for subsequent functional analysis. To demultiplex the 12 individuals pooled together for each batch, we tested both *demuxlet(Kang et al. 2018)* and a new pipeline we developed for this project (*fastdemux)* using the default parameters, resulting in cell type assignments with high agreement on genotype calls. For 165 individuals in control and LPS conditions, we had a total of 558,291 cells, with a median of 1,703 cells per sample, a median of 3,981 UMI counts, and 1,760 genes measured on average in each cell. Seurat (V5) was utilized for preprocessing, clustering, and visualizing the scRNA-seq data(Butler et al. 2018). We conducted standard preprocessing steps, including log normalization and scaling, for all counts data using Seurat’s default method. Subsequently, we performed linear dimensionality reduction on 2,000 highly variable features, resulting in 100 principal components (PCs). We also executed RunHarmony, using the library as a covariate, to correct for library-induced bias(Korsunsky et al. 2019).

To define the number of dimensions and resolution for clustering we ran clustering for several resolutions with dimension 50 (default setting) and dimension 13 (where the additional dimensions contributed little extra information as visualized in an elbow plot). At resolution 0.15, we identified 7 clusters and then performed celltyping using sctype (REF) using their immune tissue as a reference. From this, we got a score for every possible cell type and we identified 6 major PBMC cell types. For every major cell type we then identified the top assignment for the cluster in question by summing the scores of the cell types in that group. Then for each individual cell type we assigned correct assignment (1) if that individual cell type is in the top major cell type. We then calculated the sum of correct assignments vs the total to get the proportion of correct assignments for each cluster. Then for each resolution, we summed the proportion per cluster * cluster_size/total_cells so bigger clusters weighed more. Using this method, we selected dimension 13 and resolution 0.15 for downstream analysis. Uniform Manifold Approximation and Projection (UMAP) was applied to visualize the clustering results. Cells from various batches and treatments demonstrated similar clustering patterns (Figure S4-7).

### Differential gene expression analysis

For each cluster and treatment, we generated pseudobulk RNA-seq data matrix by adding together all the cells of each individual/sample resulting in a column vector where each row is a gene. We filtered out genes to keep genes expressed in 25% of the columns and removed those columns with less than 20 cells in the cluster. For some of the small clusters with less cells we may have a smaller number of individuals/columns in the pseudobulk matrix.We performed differential gene expression analysis for each variable, treatment and cell type separately using R DESeq2 package(Love, Huber, and Anders 2014). To adjust for potential confounders, we included batch, sex, age, and genotype PC1 and PC2 as covariates. To account for multiple hypothesis testing, we applied the Benjamini-Hochberg controlled FDR(Benjamini and Hochberg 1995) to a list of P-values for each run separately. DEGs were defined as those with 10% FDR. We also estimated gene expression effects of the treatments using a contrast model of LPS versus CTRL, including the same covariates as stated previously.

### Pathway enrichment analysis

Over-representation analysis (ORA) was done in ReactomePA using the Reactome pathways implemented in *enrichPathway(Yu et al. 2012)*. The ORA was independently performed for upregulated genes associated with psychological stress in each cell type and downregulated genes associated with social support in each cell type. Enrichment results were concatenated for all cell types and variables, filtered for FDR 10%, and ordered by frequency of enrichment across cell types and variables. The top 4 pathways were highlighted in Figure 3B.

### scATAC-seq raw data processing (alignment, quality control, clustering and cell type annotation)

We aligned reads for ATAC-seq data from 27 libraries by running cellranger-arc with the human reference genome (GRCh 38). Because 12 individuals were pooled together for each batch in experiments, we demultiplexed and assigned the individual identity for each cell with fastdemux. Fastdemux was selected over demuxlet due to demuxlet limitation on runtime and handling of the large number of scATAC reads. We processed these aligned reads using the ArchR(Granja et al. 2021) package with the fragment files as input data. In the ArchR, we start with the two steps (1) create arrow files including quality control, generating genome genome-wide tile matrix and gene score matrix. These large data sets are stored as HDF5-format files on disk. (2) create ArchRProject to interact with arrow files. The ArchRProject is a small size object stored in memory. For quality control, we used default settings for cellranger-arc and ArchR steps. We also kept the singlet cells with correct matching between individual and batch. A total of 400,535 cells remained in the following analysis.

For dimensional reduction, we implemented iterative latent semantic indexing (LSI) on genome-wide 500 bp tile matrix by running addIterativeLSI with default settings including the following procedures (1) compute an LSI transformation of the most accessible tiles on a small subset; (2) then project all the cells in this LSI subspace; (3) cluster cells using Seurat’s shared Nearest Neighbor approach; (4) Identify the most variable features given these defined clusters. In the second iterative, we repeat the procedure (1) and (2) to get final LSI for all the cells. In the default setting, we also excluded the LSIs that correlated with sequence depth greater than 0.75 (default value). We applied the Harmony algorithm to correct batch effects by running addHarmony with the groupBy setting to be library. We clustered cells using addClusters with the 0.07 resolution. This function implemented the graph-based clustering approach directly from Seurat. We performed Uniform Manifold Approximation and Projection(UMAP) to visualize the single cell data.

At the 0.07 resolution, we identified 10 main clusters. To recognize the immune cell types for the defined clusters, we integrated the single cell RNA-seq data from the same experiment through addGeneIntegrationMatrix. This function parallelly performed a separate alignments for the divided subsets of cells with the FindTransferAnchors from Seurat package. To bridge the features between scATAC-seq and scRNA-seq, we inferred the gene scores from chromatin accessibility using the default ArchR gene score model with the imputation of gene values by MAGIC. We transferred the RNA labels at the resolution of 0.15 to ATAC-seq data and computed the confusion matrix between the defined clusters and the transferred labels.

### Peaks calling for the defined clusters with MACS2

We performed the peak calling procedure for the 10 clusters with 0.07 resolution using MACS2 by the ArchR addReproduciblePeakSet function. It adopted an iterative overlap approach to optimize cell type-specific peaks(Corces et al. 2018). Through this procedure, we identified a robust peak set with 274,488 features. Next, we quantified the insertion counts for the new peak set with the addPeakMatrix. Lastly, we annotated the genes closest to this peak set using the hg38 reference genome with the ≤100 kb distance.

### Transcription factors motif analysis

For the robust peak set, we obtained a TF-occupied matrix for the presence of a motif in each peak using the addMotifAnnotations wrapped in the matchMotifs from motifmatchr package. Here we focused on the 692 core motifs from the JASPAR2022(Castro-Mondragon et al. 2022).

We implemented the chromVAR through the addDeviationsMatrix function to compute the TF motif activity. These values are the deviations between the sum of genome-wide chromatin accessibility (excluding sex chromosomes) of the peaks with a particular TF and the expected values. These values are also corrected for technical confounders using background peak sets(Schep et al. 2017). The z-scores (deviation/standard error) are used for downstream analysis. In order to investigate the associations between TF motif activity and psychosocial variables, we averaged the motif activity for the cells from the same individual, cell type and treatment. This resulted in a matrix for each treatment and cell type, where each row represents a motif and each column averages the motif activity across all the cells from the same individual. We then used a linear model to perform an association analysis between TF activity and a psychosocial factor (each one tested separately) and adding as confounders the top 2 genotype PCs, gender, age and batch. The differentially active motif activity (DAM) are defined as those with less than 10% FDR derived from p.adjust with Benjamini-Hochberg bonferroni correction.

### Motif enrichment analysis

We derived the TF-regulated-gene sets for each individual TF with the following criteria, (1) the peaks with the binding of TF which is from TF-occupied matrix. For each cell type, we considered the peaks measured at least 1% cells. (2) the genes closest to these TF peak sets. We performed two types of TF motif enrichment analysis for DEGs in the corresponding cell types and psychosocial factors using Fisher’s exact test, (1) Perform an enrichment test for individual TF to identify what individual TFs are more important in the regulation of DEGs; (2) Perform an overall test to examine if DEGs are enriched in DAMs.

## Acknowledgments

We thank members of the Luca/Pique-Regi/Zilioli groups and the CZI African Ancestry Network groups for helpful discussions. This work was funded by R01GM109215 (FL, RPR), R01HL153377 (SZ), P30AG015281 (SZ), P50MD017351 (SZ), P30ES020957 Pilot grant (FL), CZI Ancestry Networks (FL, RPR). AI tools were used to edit portions of the initial draft of the manuscript.

## Notes

### Competing Interest Statement

The authors have declared no competing interest.

