## Supplemental material for "Psychological stress and social support are associated with opposing single-cell pro-inflammatory gene regulatory mechanisms in adults"

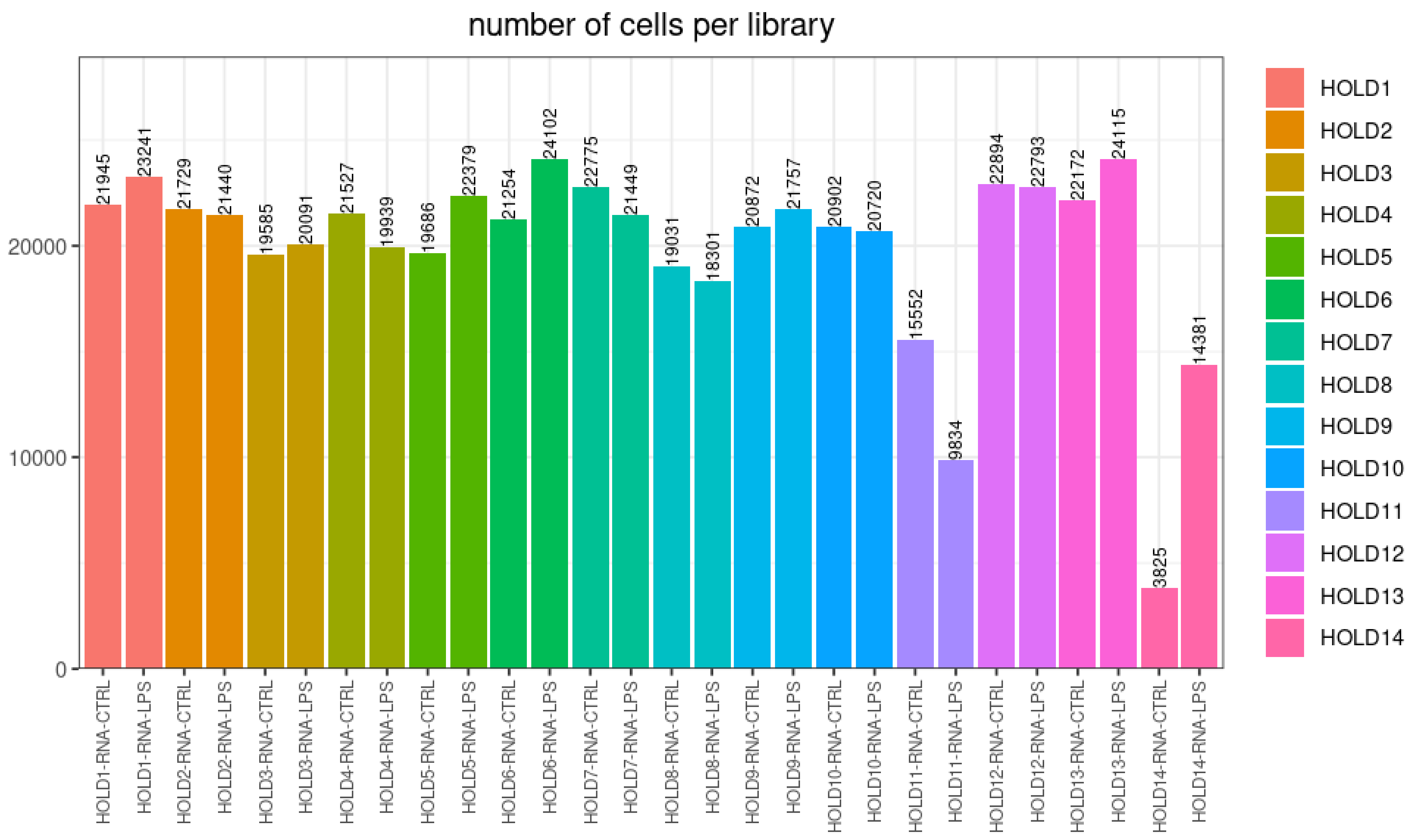


**Figure S1.** Number of cells per single cell library

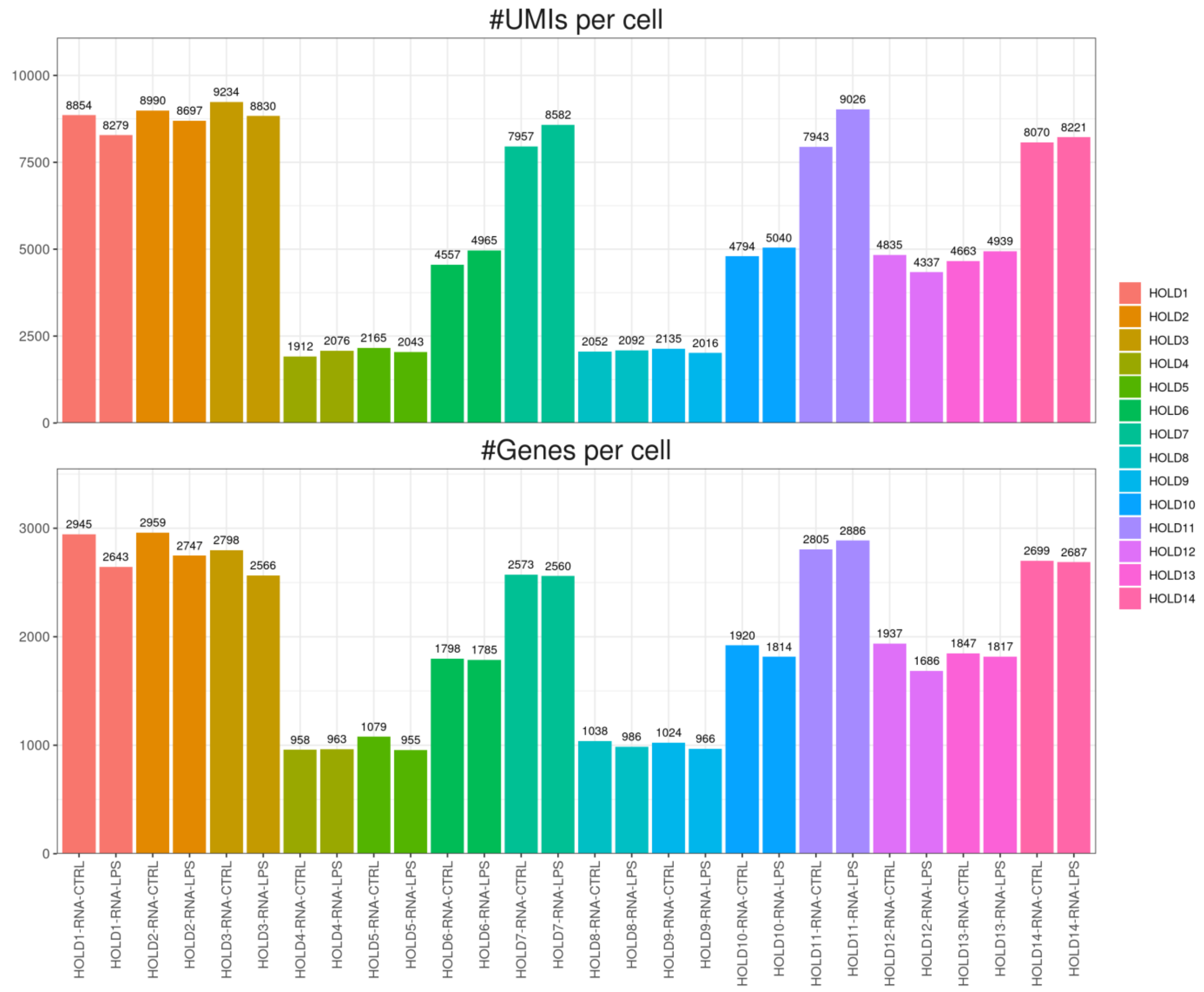


**Figure S2.** Number of unique molecular identifiers (UMIs) and gene features per cell per library

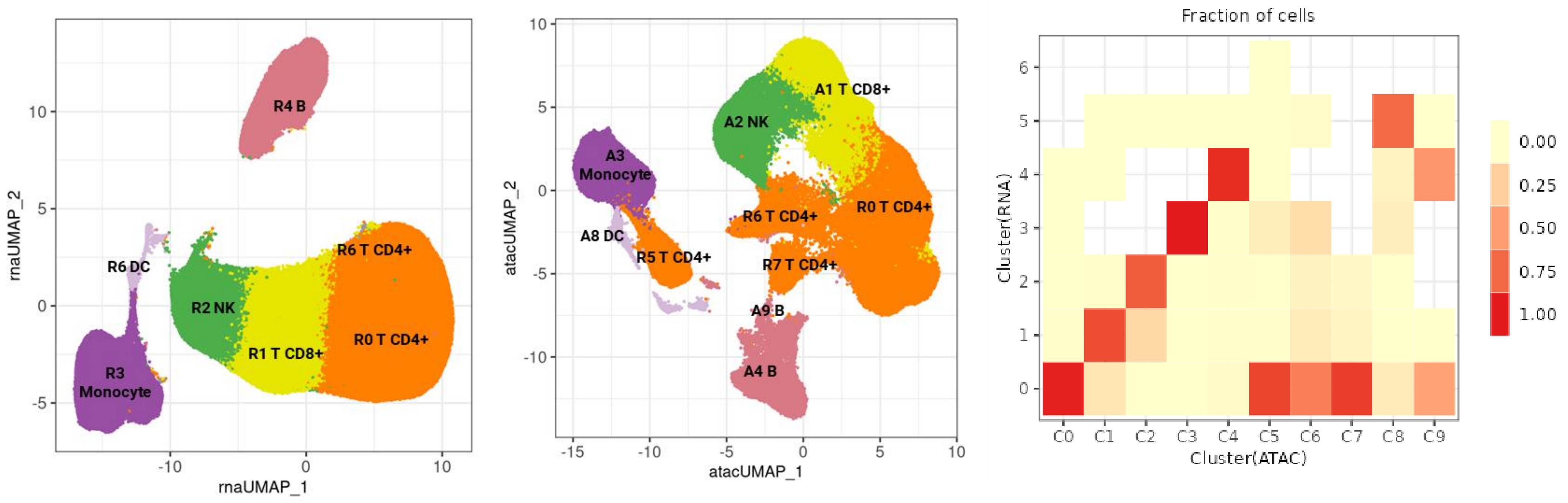


**Figure S3.** UMAP of annotated PBMCs from scRNA-seq and scATAC-seq

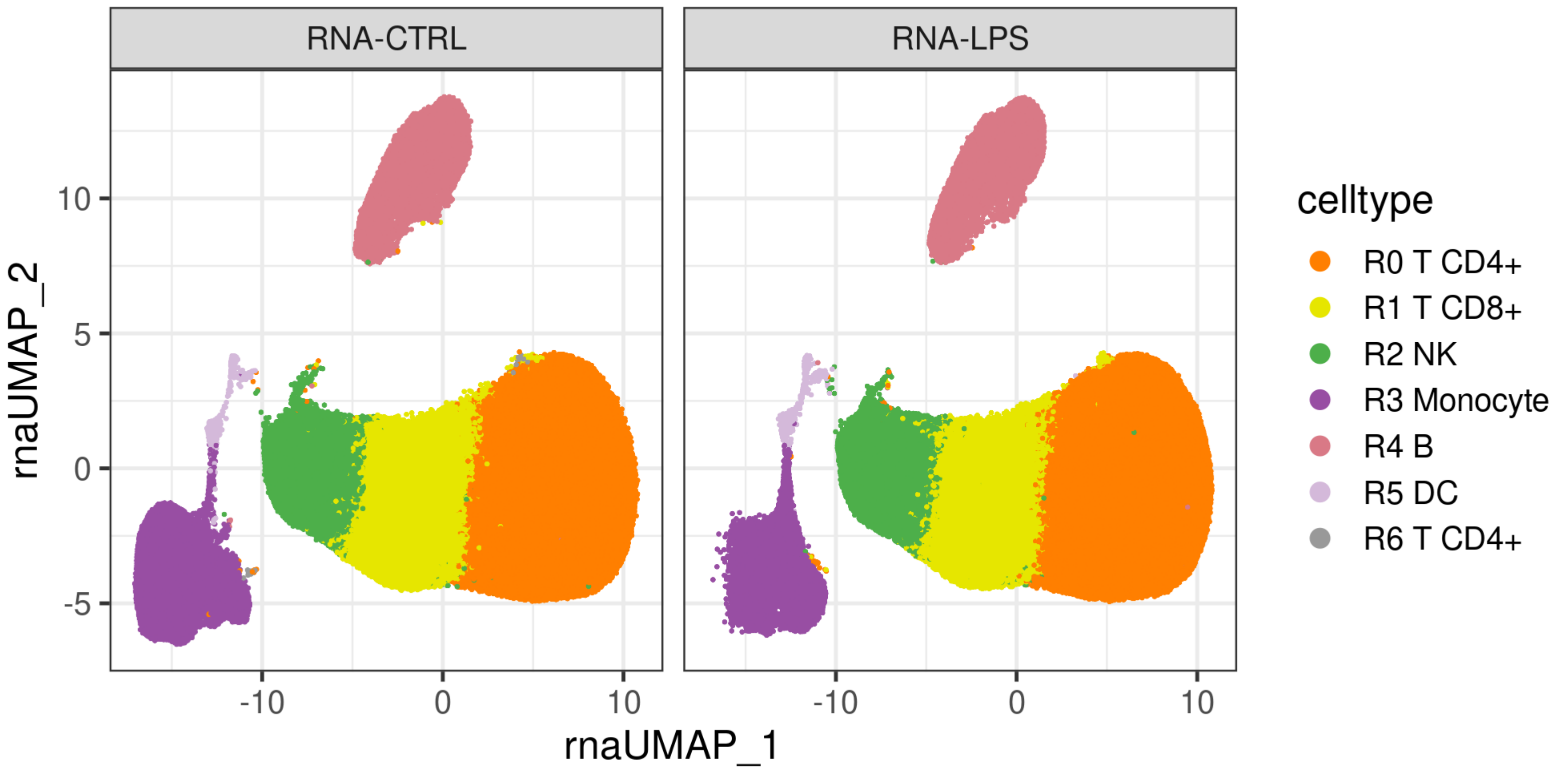


**Figure S4.** scRNA-seq UMAP split by treatment

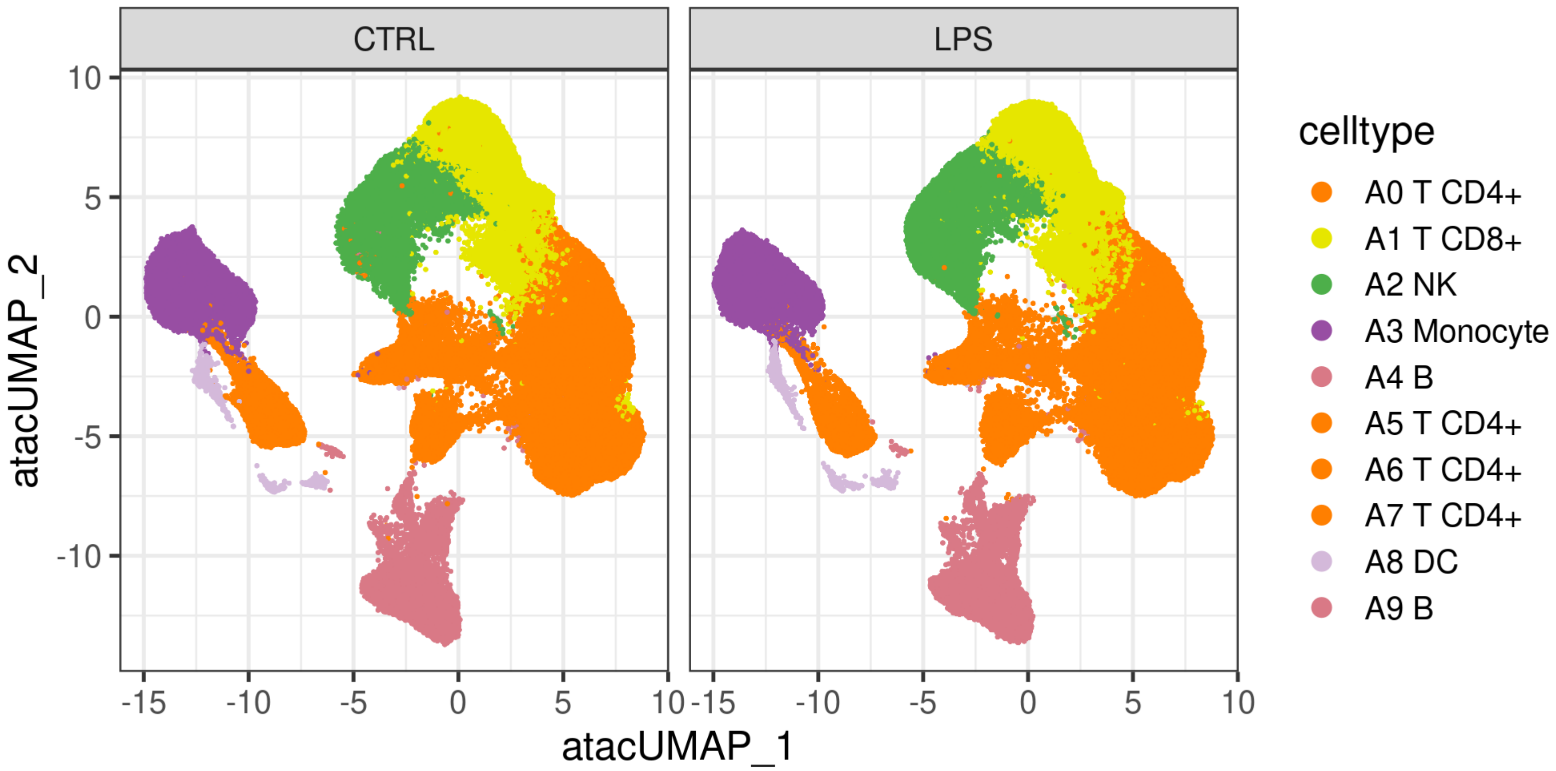


**Figure S5.** scATAC-seq UMAP split by treatment

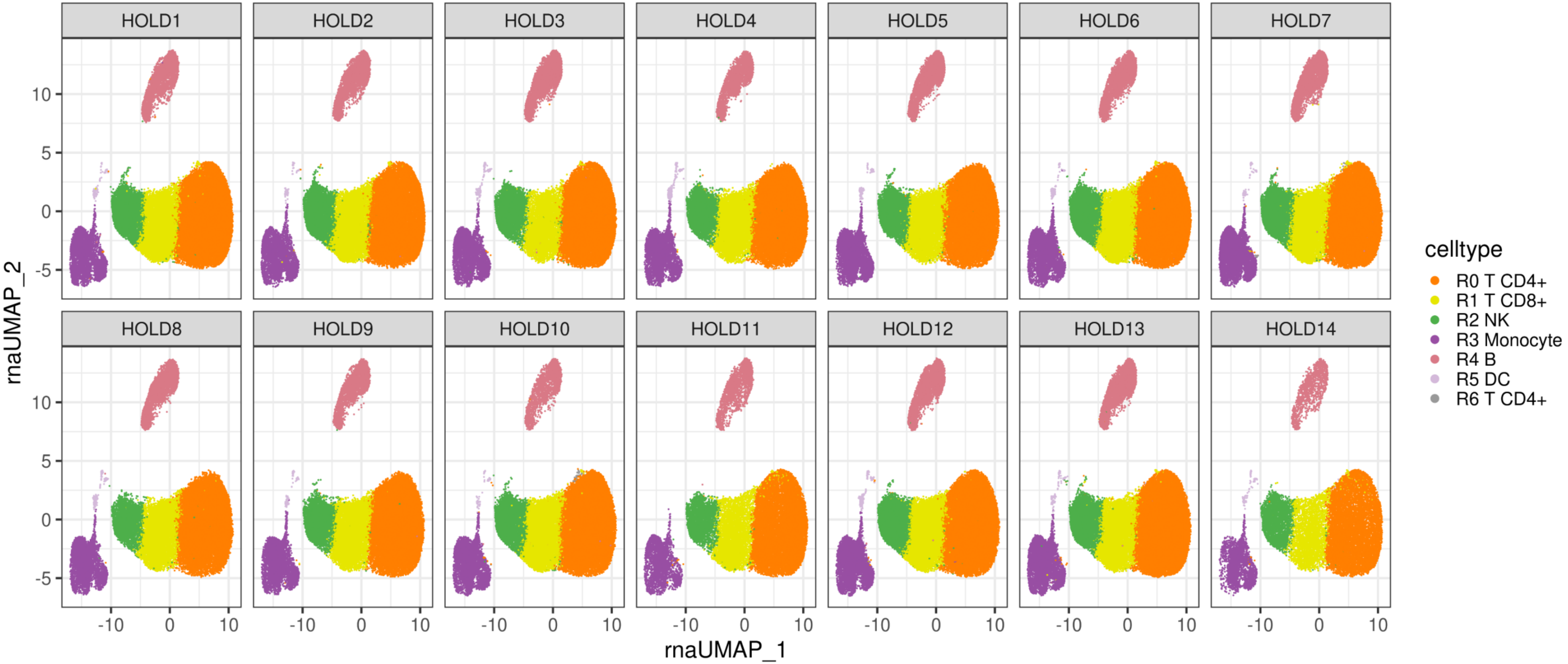


**Figure S6.** scRNA-seq UMAP split by batch

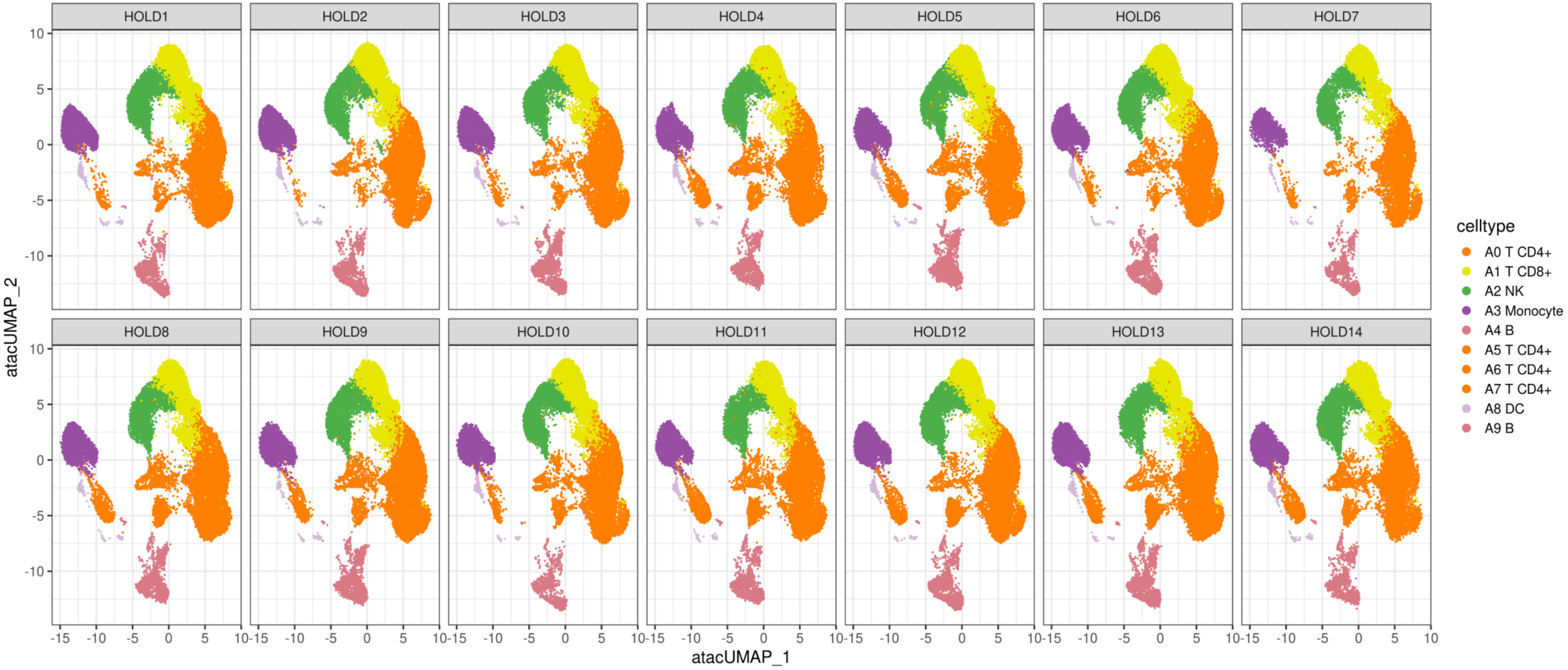


**Figure S7.** scATAC-seq UMAP split by batch

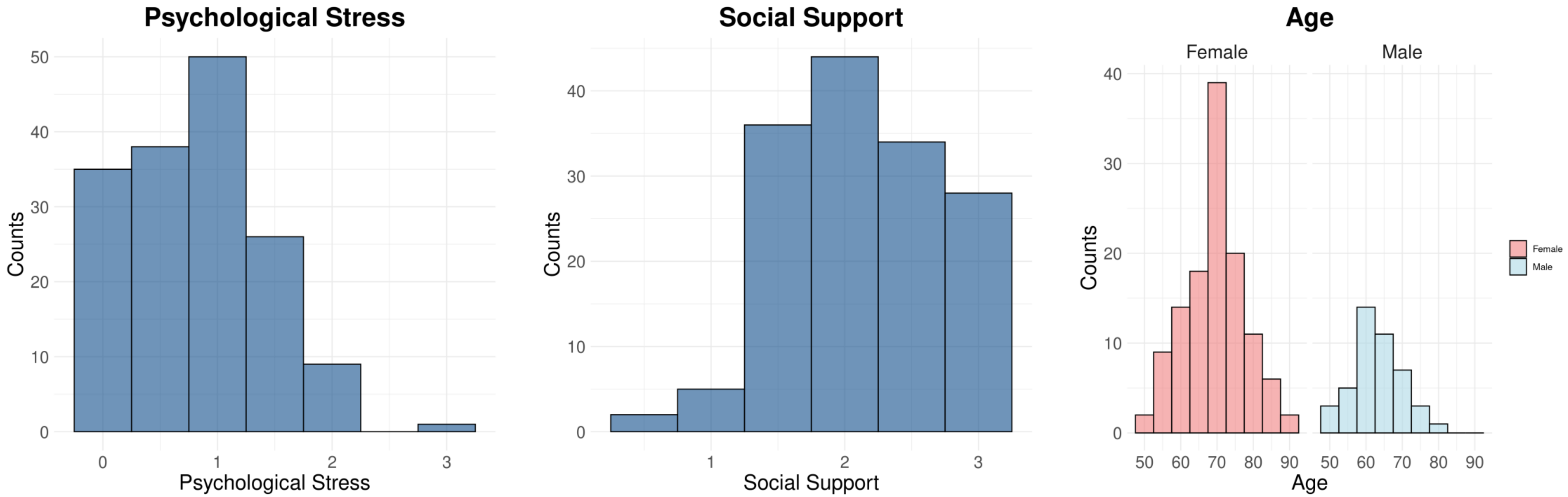


**Figure S8.** Distribution of (A) psychological stress scores (B) social support scores (C) age separated by sex

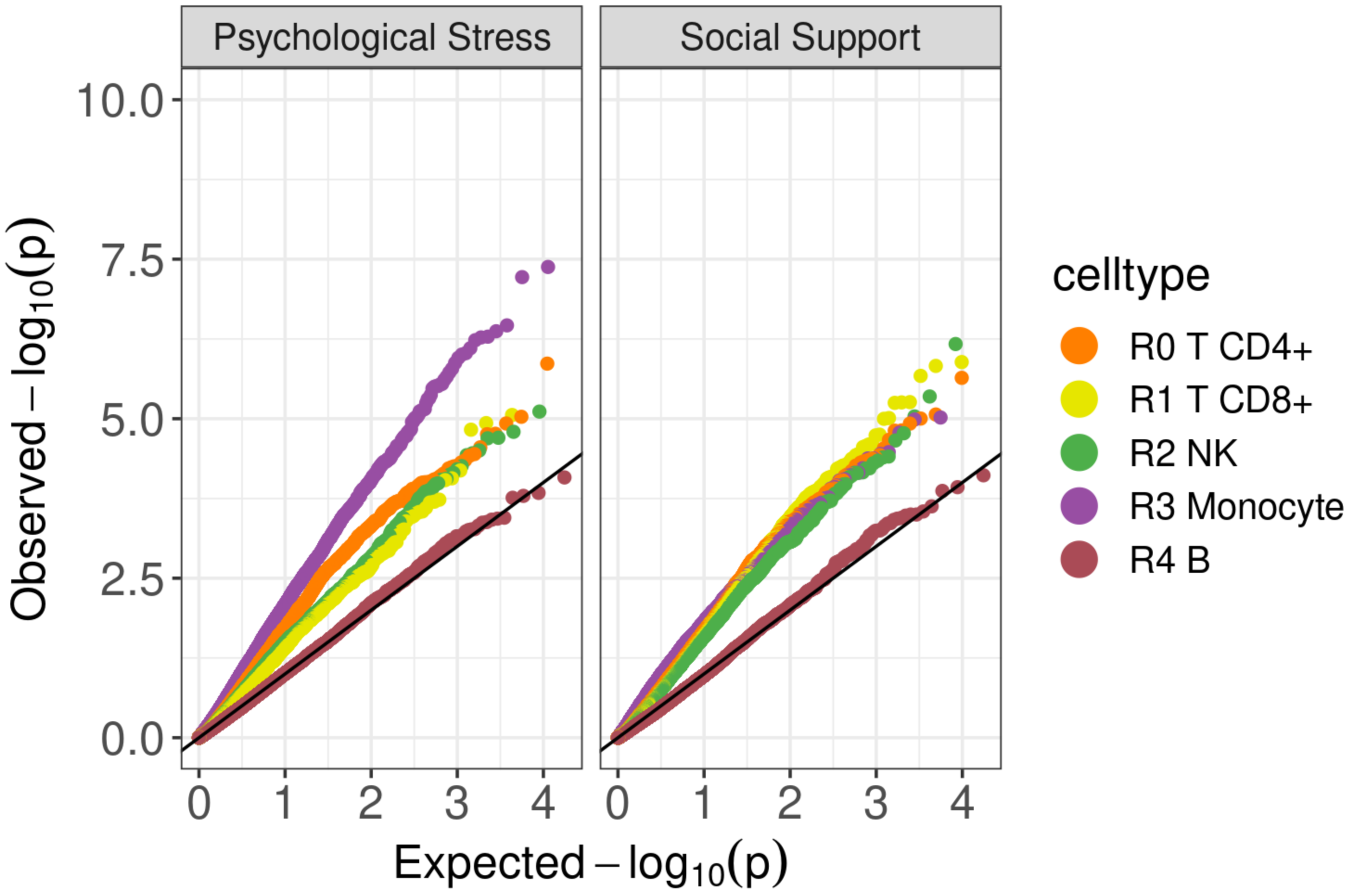


**Figure S9.** Q-Q plots of differentially expressed genes associated with psychological stress and social support (colored by different immune cell types), x-axis representing expected –log10 p values and y-axis representing observed –log10 p values.

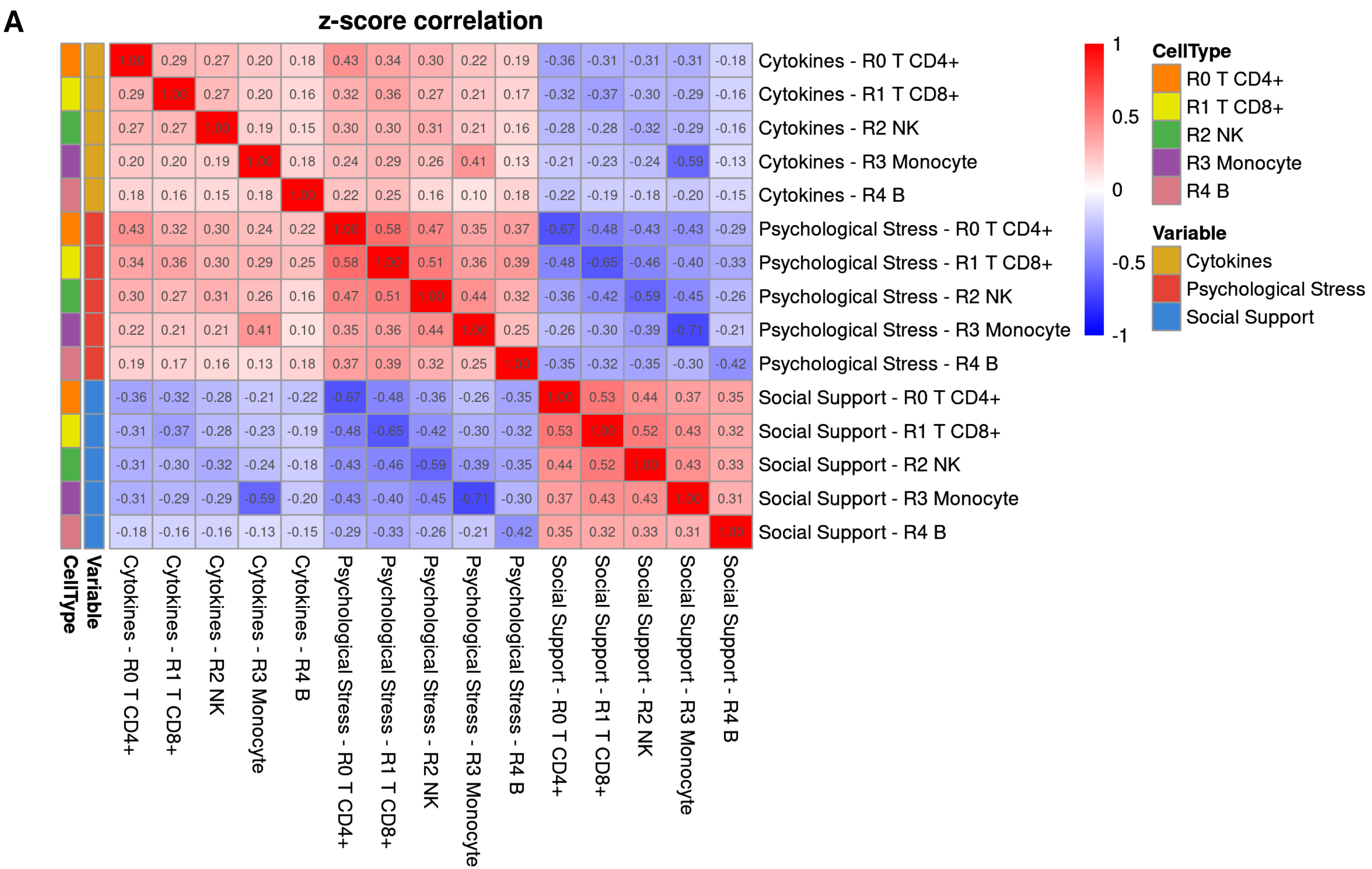


**Figure S10.** Correlation of global gene expression effect size cytokines with social support and perceived stress. (A) Heatmap of Pearson's correlation of z-scores of cytokine gene expression effect with psychological stress and social support for each cell type. All significant values p-values 0.05.

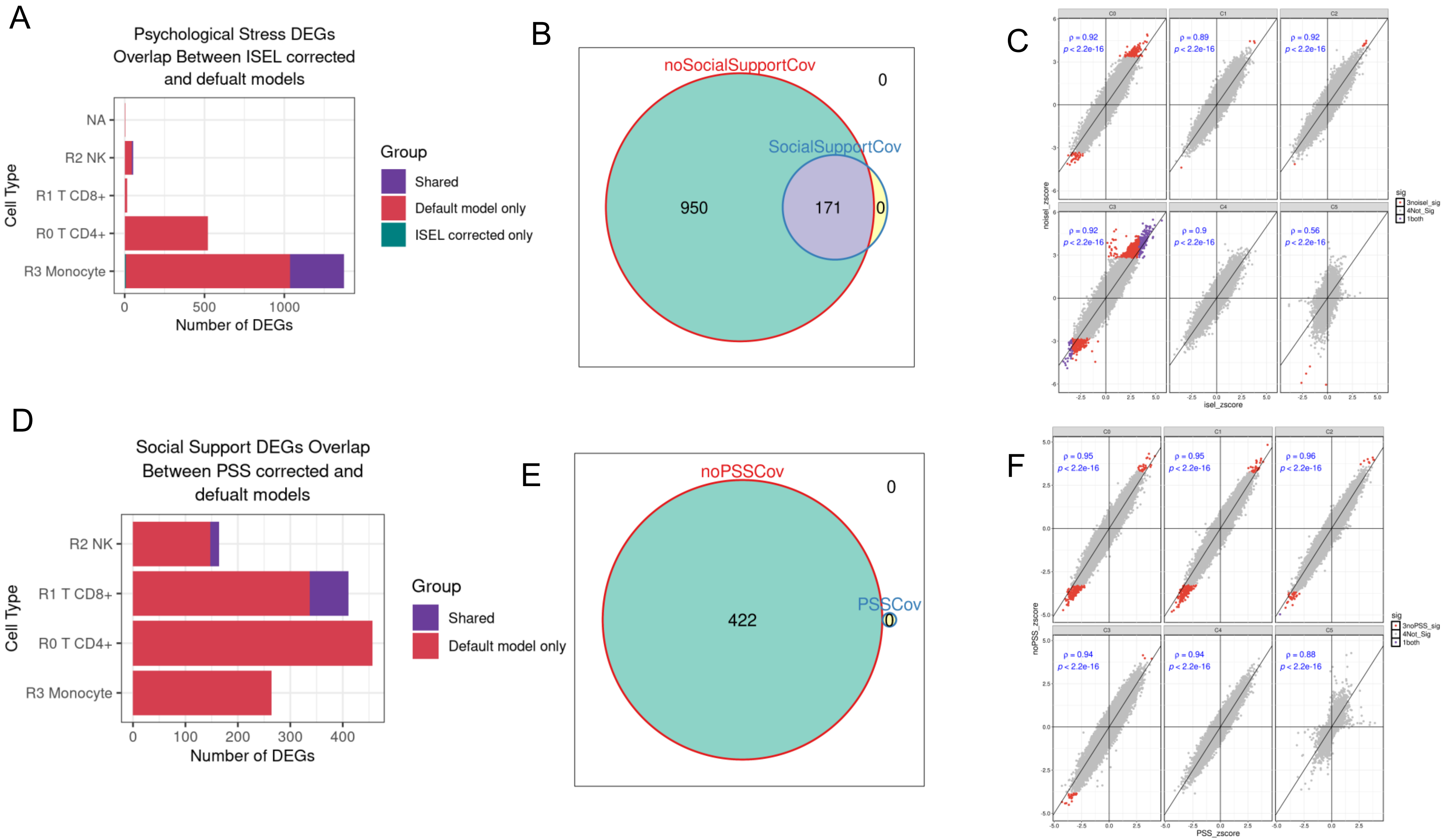


**Figure S11.** Psychological stress associated gene expression changes after controlling for social support (A) Bar plot showing the shared and unique number of psychological stress associated DEGs in each gene expression model. (B) Scatter plot of z-score correlation for all tested genes between social support corrected model and default model, colored by significance at 10% FDR (grey for significant, blue for social support corrected, red for default model, purple for both models). (C) Venn diagram of shared and unique number of DEGs across all clusters for social support corrected and default model. ). Default GE model: GE ~ genPC1 + genPC2 + sex + age + PSS. Social support corrected GE model: GE ~ genPC1 + genPC2 + sex + age + ISEL + PSS. Genes tested are fixed between the two models.

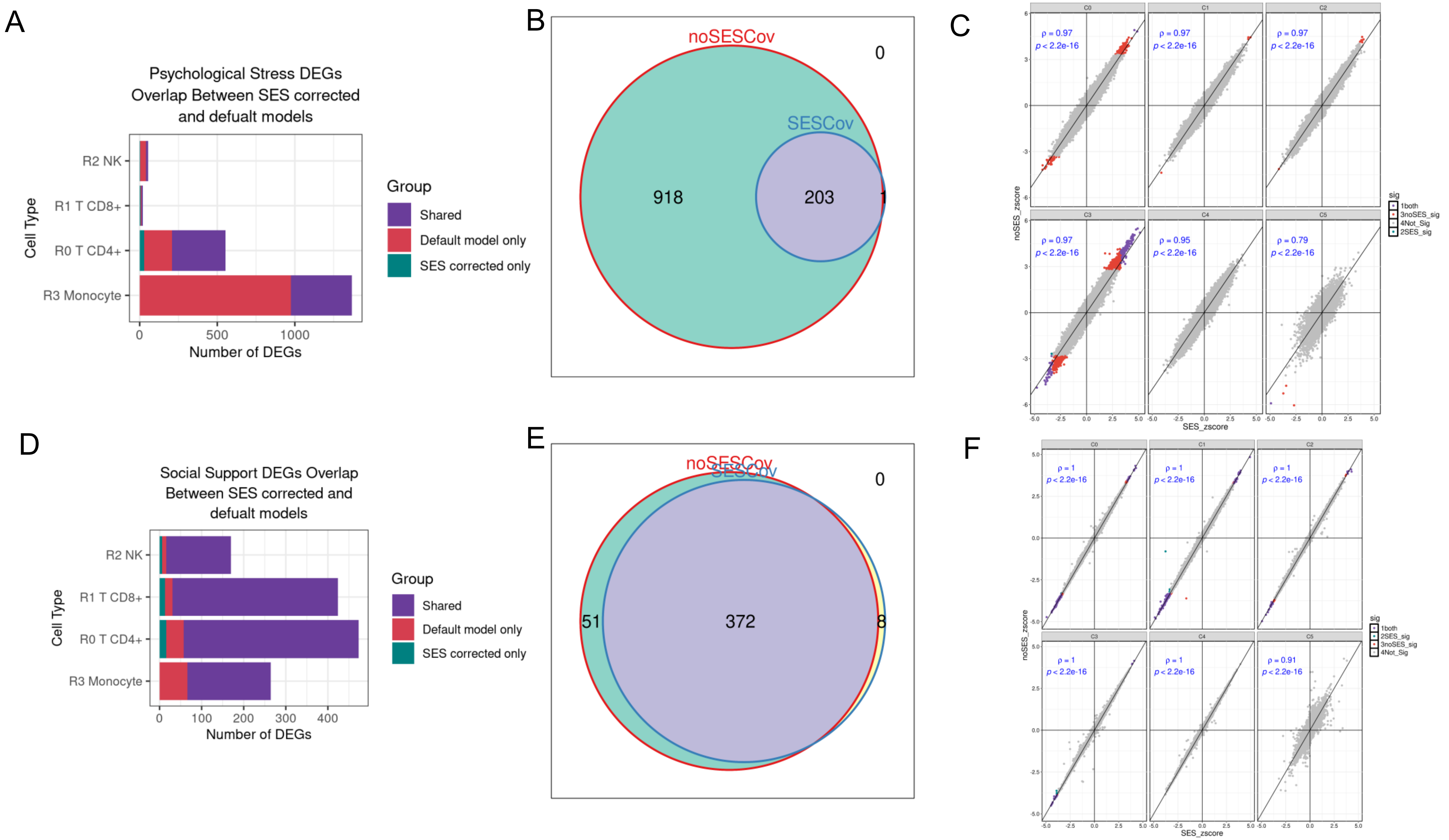


**Figure S12.** Psychological stress (top row) and social support (bottom row) associated gene expression changes after controlling for socio economic status (SES). (A and D) Bar plot showing the shared and unique number DEGs in each gene expression model. (B and E) Venn diagram of shared and unique number of DEGs across all clusters for SES corrected model and default model (C and F) Scatter plot of z-score correlation for all tested genes between SES corrected model and default model, colored by significance at 10% FDR (grey for significant, blue for SES corrected, red for default model, purple for both models). Default GE model: GE ~ genPC1 + genPC2 + sex + age + ISEL or PSS. SES corrected GE model: GE ~ genPC1 + genPC2 + sex + age + SES + ISEL or PSS.

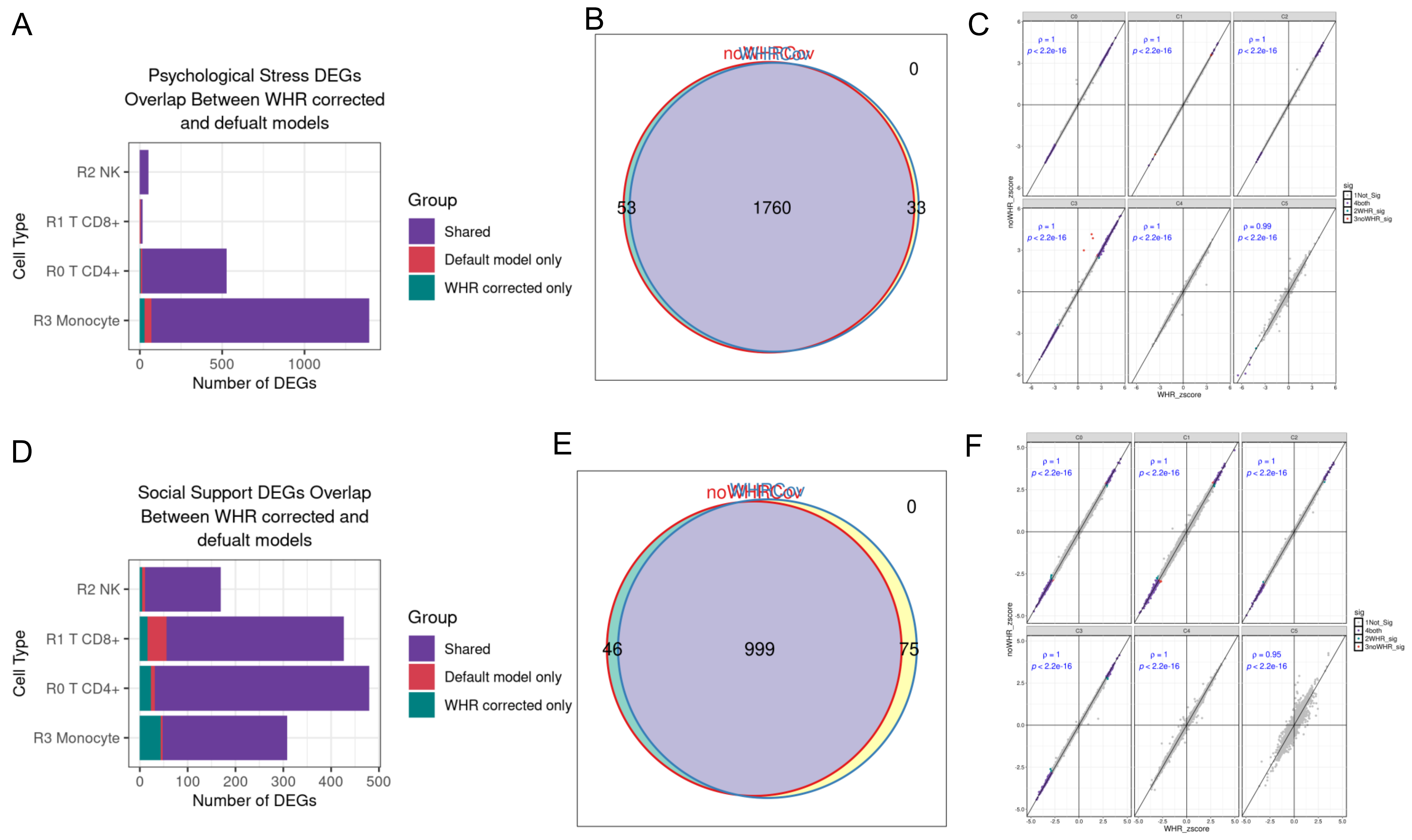


**Figure S13.** Psychological stress (top row) and social support (bottom row) associated gene expression changes after controlling for waist-to-hip ratio (WHR). (A and D) Bar plot showing the shared and unique number DEGs in each gene expression model. (B and E) Venn diagram of shared and unique number of DEGs across all clusters for WHR corrected and default model. (C and F) Scatter plot of z-score correlation for all tested genes between WHR corrected model and default model, colored by significance at 10% FDR (grey for significant, blue for WHR corrected, red for default model, purple for both models. Default GE model: GE ~ genPC1 + genPC2 + sex + age + ISEL or PSS. WHR corrected GE model: GE ~ genPC1 + genPC2 + sex + age + WHR + ISEL or PSS.

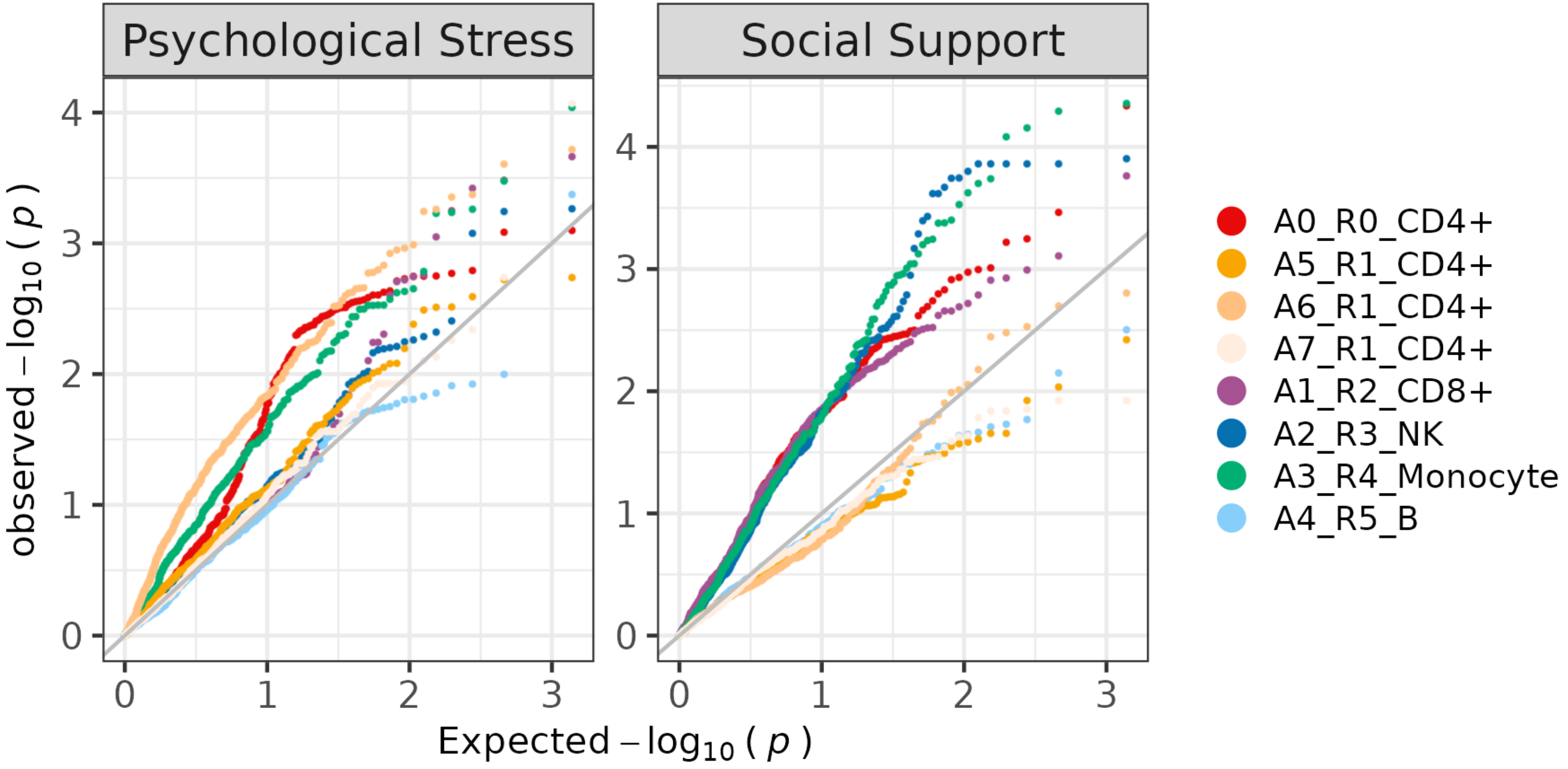


**Figure S14.** Q-Q plots of TF motifs activity associated with psychosocial factors (colored by different immune cell types), x-axis representing expected –log10 p values and y-axis representing observed –log10 p values.

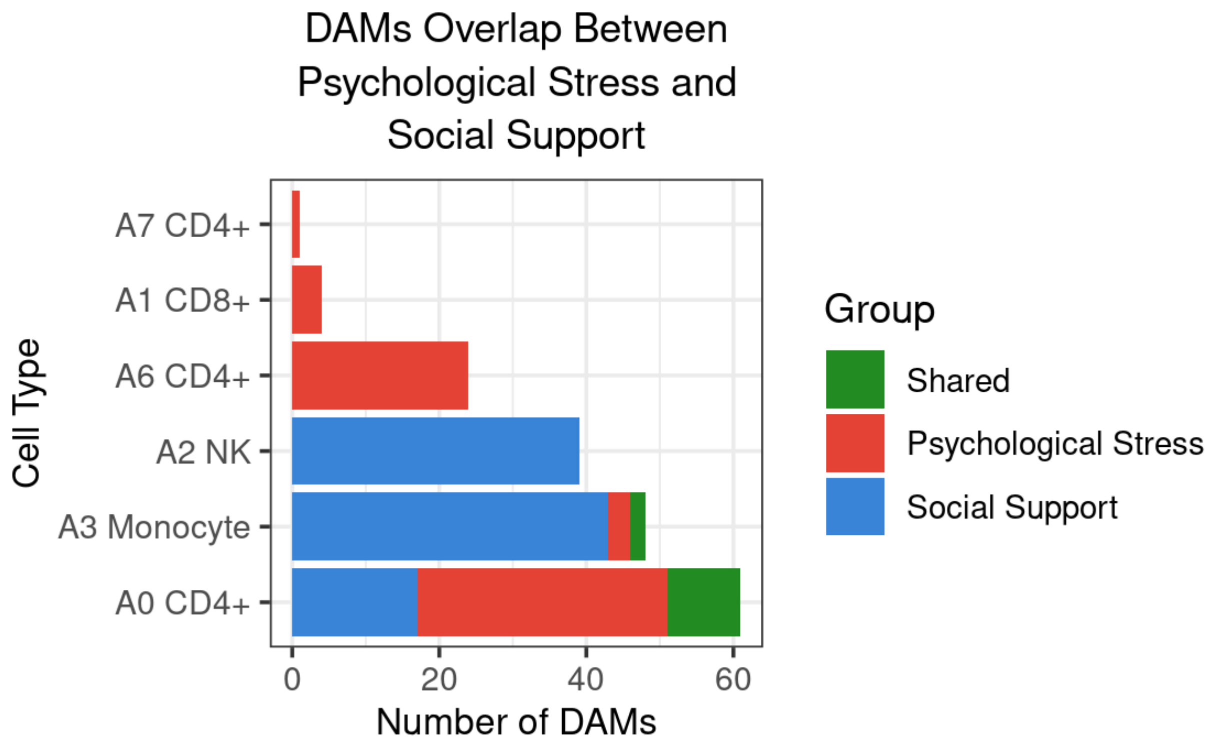


**Figure S15.** Number of shared and unique DAMs in psychological stress and social support per cell type

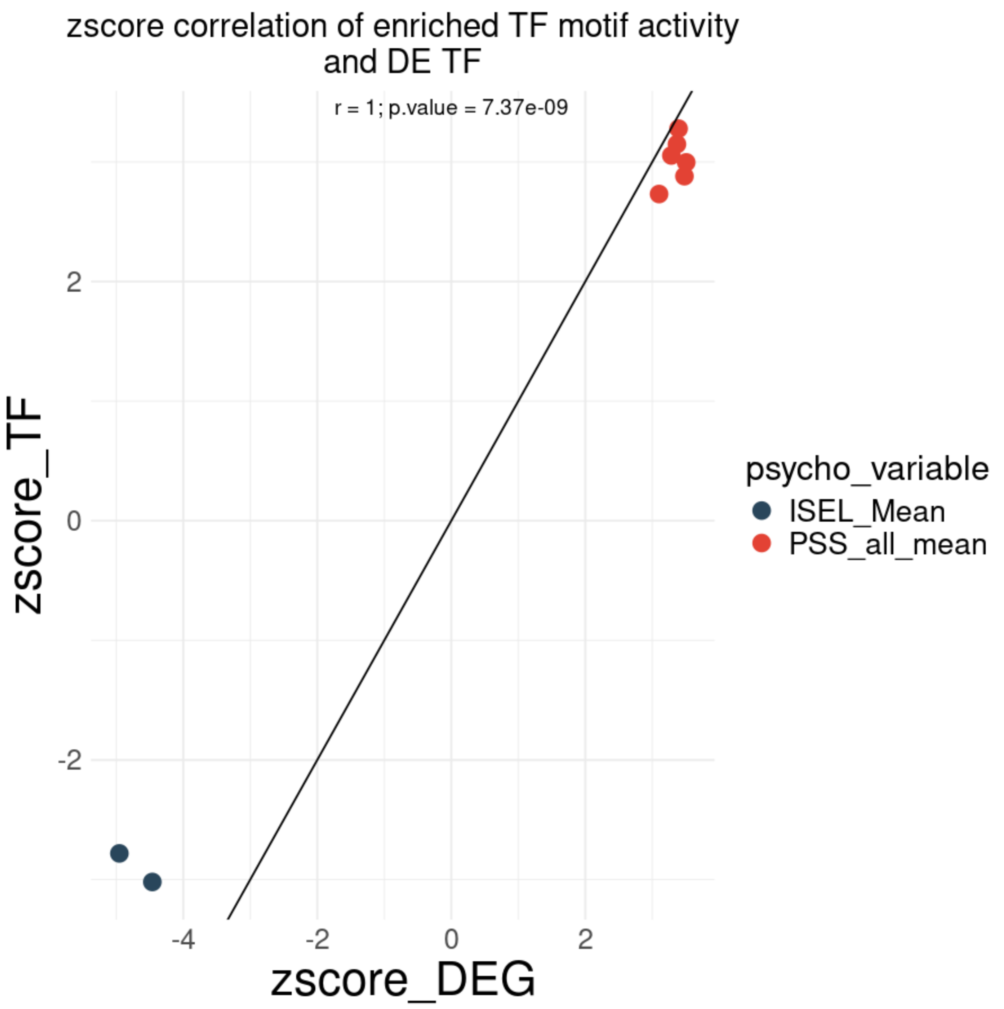


**Supplementary Figure S16.** z-score correlation of changes in motif activity and TF gene expression (DEG) associated with perceived stress (red) and social support (blue) in CD4+ T cells.

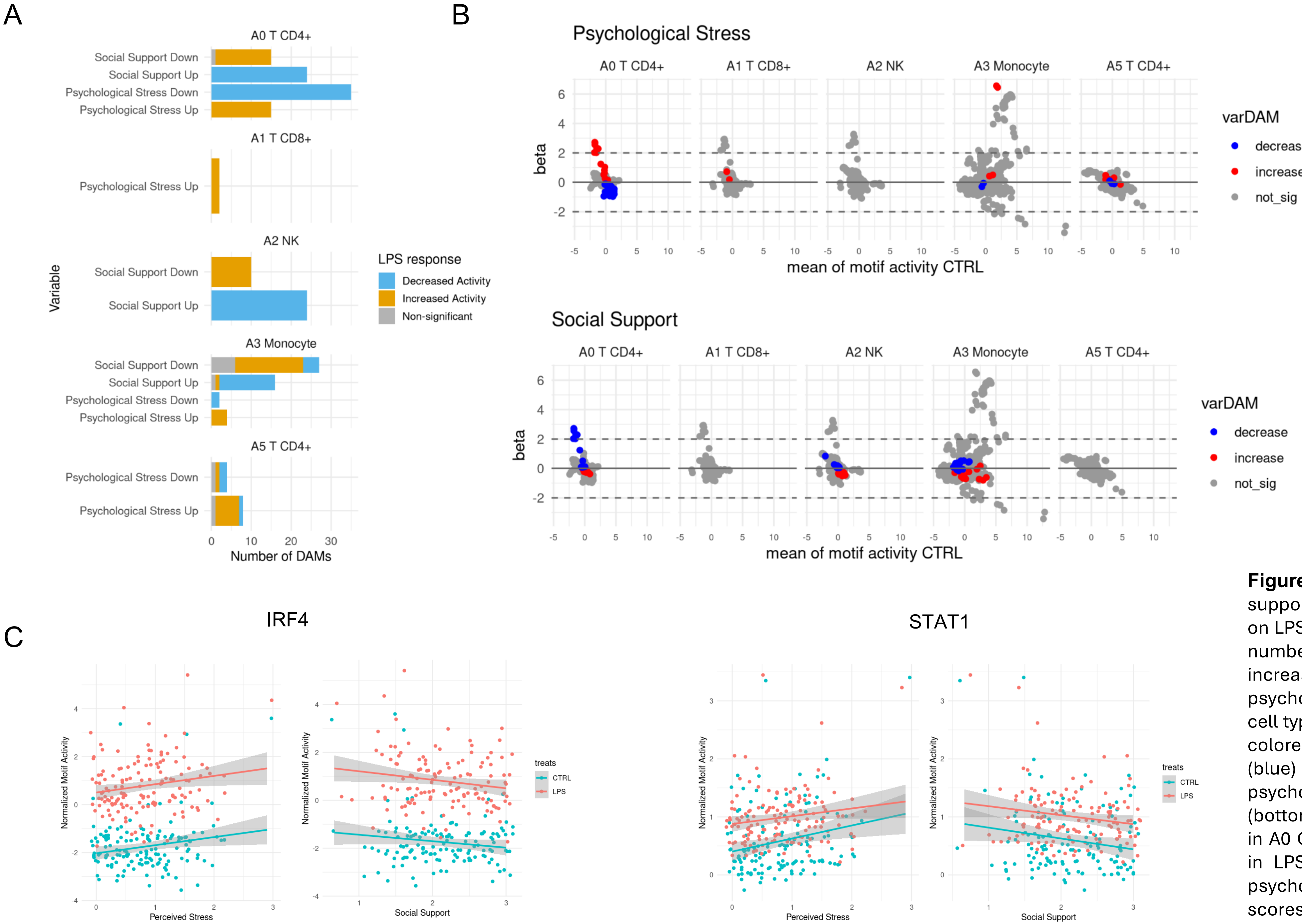


**Figure S17.** Psychological stress and social support associated differentially motif activity on LPS treated cells (A) Stacked bar plot of the number of LPS response DAMs that have increased or decreased motif activity with psychological stress and social support, per cell type. (B) MA plot of LPS vs CTRL treatment colored by increased (red) and decreased (blue) motif activity associated with psychological stress (top) and social support (bottom). (C) Normalized motif activity of IRF4 in A0 CD4+ T cell and STAT1 in A3 monocytes in LPS and CTRL treatment correlated with psychosocial stress and social support scores.

**Table S1.** Interpersonal Support Evaluation (ISEL) Questionnaire

| | |
|---|---|
| If I wanted to go on a trip for a day (for example, to the country or mountains), I would have a hard time finding someone to go with me | ISEL_01 |
| I feel that there is no one I can share my most private worries and fears with | ISEL_02 |
| If I were sick, I could easily find someone to help me with my daily chores | ISEL_03 |
| There is someone I can turn to for advice about handling problems with my family | ISEL_04 |
| If I decided one afternoon that I would like to go to a movie that evening, I could easily find someone to go with me | ISEL_05 |
| When I need suggestions on how to deal with a personal problem, I know someone I can turn to | ISEL_06 |
| I don't often get invited to do things with others | ISEL_07 |
| If I had to go out of town for a few weeks, it would be difficult to find someone who would look after my house or apartment (plants, pets, garden, etc.) | ISEL_08 |
| If I wanted to have lunch with someone, I could easily find someone to join me | ISEL_09 |
| If I was stranded 10 miles from home, there is someone I could call who could come and get me | ISEL_10 |
| If a family crisis arose, it would be difficult to find someone who could give me good advice about how to handle it | ISEL_11 |
| If I needed some help in moving to a new house or apartment, I would have a hard time finding someone to help me | ISEL_12 |

**Table S2.** Perceived psychological stress (PSS) questionnaire. Averaged over 5 day.

| **Your Experiences Today** | | |
|---|---|---|
| Thinking about today, how often did you feel that you were unable to control the important things of the day? | 1 *(never)*<br>2 *(almost never)*<br>3 *(sometimes)*<br>4 *(fairly often)*<br>5 *(very often)* | Exp01 |
| Thinking about today, how often did you feel confident about your ability to handle your personal problems? | 1 *(never)*<br>2 *(almost never)*<br>3 *(sometimes)*<br>4 *(fairly often)*<br>5 *(very often)* | Exp02 |
| Thinking about today, how often did you feel that things were going your way? | 1 *(never)*<br>2 *(almost never)*<br>3 *(sometimes)*<br>4 *(fairly often)*<br>5 *(very often)* | Exp03 |
| Thinking about today, how often did you feel that difficulties were piling up so high that you could not overcome them? | 1 *(never)*<br>2 *(almost never)*<br>3 *(sometimes)*<br>4 *(fairly often)*<br>5 *(very often)* | Exp04 |

**Table S3.** scRNA cluster stats

| celltype | count | percentage |
|---|---|---|
| R0 T CD4+ | 274206 | 49.12 |
| R1 T CD8+ | 105539 | 18.9 |
| R2 NK | 67129 | 12.02 |
| R3 Monocyte | 65652 | 11.76 |
| R4 B | 44198 | 7.92 |
| R5 DC | 1495 | 0.27 |
| R6 T CD4+ | 72 | 0.01 |

**Table S4.** scATAC cluster stats

| celltype | count | percentage |
|---|---|---|
| A0 T CD4+ | 155930 | 38.93 |
| A1 T CD8+ | 77321 | 19.3 |
| A2 NK | 64814 | 16.18 |
| A3 Monocyte | 45315 | 11.31 |
| A4 B | 26241 | 6.55 |
| A5 T CD4+ | 9814 | 2.45 |
| A6 T CD4+ | 9633 | 2.41 |
| A7 T CD4+ | 7481 | 1.87 |
| A8 DC | 2524 | 0.63 |
| A9 B | 1462 | 0.37 |

**Table S5.** Psychological stress associated DEGs after correcting for Social support compared to the default model

| shared | Default_only | ISELcov_only | total_Default_DEGs | percent_shared | percent_Default_only |
|---|---|---|---|---|---|
| 347 | 1609 | 9 | 1956 | 17.7 | 82.3 |

**Table S6.** Social Support associated DEGs after correcting for psychological stress compared to the default model

| shared | Default_only | PSScov_only | total_Default_DEGs | percent_shared | percent_Default_only |
|---|---|---|---|---|---|
| 91 | 1205 | 0 | 1296 | 7 | 93 |

**Table S7.** Number and percentage of DEGs after correcting for SES compared to the default model

| Variable | shared | Default_only | SEScov_only | total_Default_DEGs | percent_shared | percent_Default_only |
|---|---|---|---|---|---|---|
| Psychological Stress | 759 | 1197 | 38 | 1956 | 38.8 | 61.2 |
| Social Support | 1161 | 135 | 35 | 1296 | 89.6 | 10.4 |

**Table S8.** Number and percentage of DEGs after correcting for WHR compared to the default model

| Variable | shared | Default_only | WHRcov_only | total_Default_DEGs | percent_shared | percent_Default_only |
|---|---|---|---|---|---|---|
| Psychological Stress | 1896 | 60 | 32 | 1956 | 96.9 | 3.1 |
| Social Support | 1238 | 58 | 88 | 1296 | 95.5 | 4.5 |

**Table S9.** Break down by up and down regulated genes in ISEL/PSS and LPS

| celltype | ISEL_down_LPS_down | ISEL_down_LPS_up | ISEL_up_LPS_down | ISEL_up_LPS_up | PSS_down_LPS_down | PSS_down_LPS_up | PSS_up_LPS_down | PSS_up_LPS_up |
|---|---|---|---|---|---|---|---|---|
| R0 T CD4+ | 2 | 223 | 35 | 2 | 60 | 2 | 3 | 199 |
| R1 T CD8+ | 1 | 238 | 55 | 3 | 0 | 0 | 0 | 10 |
| R2 NK | 1 | 107 | 21 | 0 | 6 | 2 | 2 | 24 |
| R3 Monocyte | 20 | 80 | 26 | 4 | 376 | 22 | 51 | 510 |
| total | 24 | 648 | 137 | 9 | 442 | 26 | 56 | 743 |

**Table S10.** Break down by up and down motif activity in ISEL/PSS and LPS

| celltype | ISEL_down_LPS_down | ISEL_down_LPS_up | ISEL_up_LPS_down | ISEL_up_LPS_up | PSS_down_LPS_down | PSS_down_LPS_up | PSS_up_LPS_down | PSS_up_LPS_up |
|---|---|---|---|---|---|---|---|---|
| A0 T CD4+ | 0 | 14 | 24 | 0 | 35 | 0 | 0 | 15 |
| A1 T CD8+ | 0 | 0 | 0 | 0 | 0 | 0 | 0 | 2 |
| A2 NK | 0 | 10 | 24 | 0 | 0 | 0 | 0 | 0 |
| A3 Monocyte | 4 | 17 | 14 | 1 | 2 | 0 | 0 | 4 |
| A5 T CD4+ | 0 | 0 | 0 | 0 | 2 | 1 | 1 | 6 |
| total | 4 | 41 | 62 | 1 | 39 | 1 | 1 | 27 |

**Table S11.** Variable distribution statistics

| | Min | Max | Median | Mean | SD | Variance |
|---|---|---|---|---|---|---|
| Psycholgoical Stress | 0.00 | 2.90 | 0.80 | 0.81 | 0.56 | 0.31 |
| Social support | 0.67 | 3.00 | 2.17 | 2.16 | 0.58 | 0.34 |
| Cytokines | -1.73 | 1.79 | -0.08 | -0.02 | 0.69 | 0.47 |
